# Deep neural network models of sound localization reveal how perception is adapted to real-world environments

**DOI:** 10.1101/2020.07.21.214486

**Authors:** Andrew Francl, Josh H. McDermott

## Abstract

Mammals localize sounds using information from their two ears. Localization in real-world conditions is challenging, as echoes provide erroneous information, and noises mask parts of target sounds. To better understand real-world localization we equipped a deep neural network with human ears and trained it to localize sounds in a virtual environment. The resulting model localized accurately in realistic conditions with noise and reverberation, outperforming alternative systems that lacked human ears. In simulated experiments, the network exhibited many features of human spatial hearing: sensitivity to monaural spectral cues and interaural time and level differences, integration across frequency, and biases for sound onsets. But when trained in unnatural environments without either reverberation, noise, or natural sounds, these performance characteristics deviated from those of humans. The results show how biological hearing is adapted to the challenges of real-world environments and illustrate how artificial neural networks can extend traditional ideal observer models to real-world domains.

## Introduction

Why do we see or hear the way we do? Perception is believed to be adapted to the world – shaped over evolution and development to help us survive in our ecological niche. Yet adaptedness is often difficult to test. Many phenomena are not obviously a consequence of adaptation to the environment, and perceptual traits are often proposed to reflect implementation constraints rather than the consequences of performing a task well. Well-known phenomena attributed to implementation constraints include aftereffects (1, 2), masking (3, 4), poor visual motion and form perception for equiluminant color stimuli (5), and limits on the information that can be extracted from high-frequency sound (6–8).

Evolution and development can be viewed as an optimization process that produces a system that functions well in its environment. The consequences of such optimization for perceptual systems have traditionally been revealed by ideal observer models – systems that perform a task optimally under environmental constraints (9), and whose behavioral characteristics can be compared to actual behavior. Ideal observers are typically derived analytically, but as a result are often limited to simple psychophysical tasks (10–13). Despite recent advances, such models remain intractable for many real-world behaviors. Rigorously evaluating adaptedness has thus remained out of reach for many domains.

Sound localization is one domain of perception where the relationship of behavior to environmental constraints has not been straightforward to evaluate. The basic outlines of spatial hearing have been understood for decades (14–17). Time and level differences in the sound that enters the two ears provide cues to a sound’s location, and location-specific filtering by the ears, head, and torso provide monaural cues that help resolve ambiguities in binaural cues (Figure 1A). However, in real-world conditions, background noise masks or corrupts cues from sources to be localized, and reflections provide erroneous cues to direction (18). Classical models based on these cues thus cannot replicate real-world localization behavior (19–21). Instead, modeling efforts have focused on accounting for observed neuronal tuning in early stages of the auditory system rather than behavior (22–27), or have modeled behavior in simplified experimental conditions using particular cues (21, 28–31). Engineering systems must solve localization in real-world conditions, but typically adopt approaches that diverge from biology, using more than two microphones and/or not leveraging ear/head filtering (32–34). As a result we lack quantitative models of how biological organisms localize sounds in realistic conditions.

**Figure 1.**
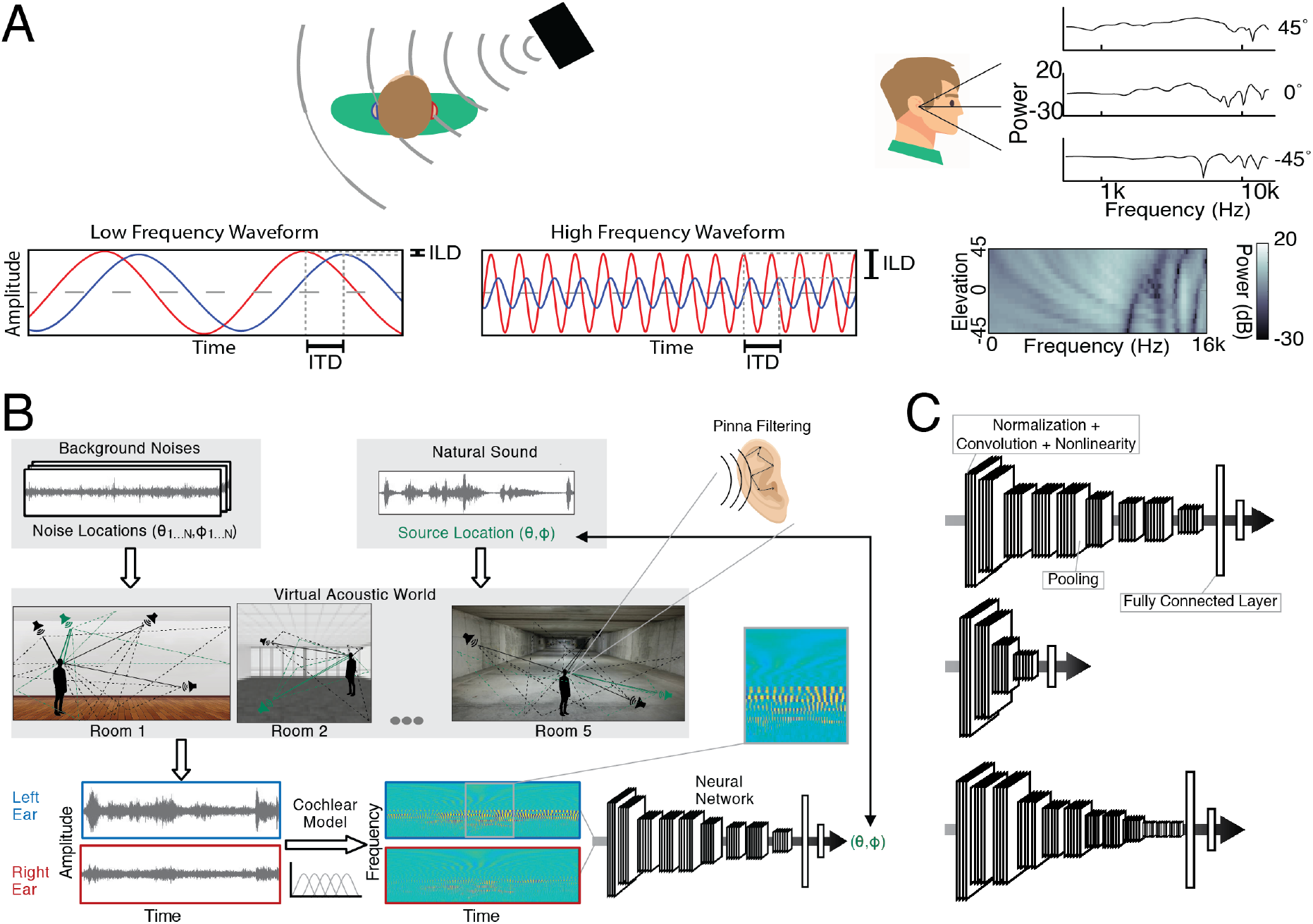
Overview of spatial hearing and training procedure. A. Schematic of cues to sound location available to human listeners: interaural time and level differences (left and center) and spectral differences (right). Time and level differences are shown for a low and high frequency sinusoids (left and center, respectively). Note that the level difference is small for the low frequency, and that the time difference is ambiguous for the high frequency. B. Training procedure. Natural sounds (in green) were rendered at a location in a room, with noises (natural sound textures, in black) placed at other locations. The rendering included the direction-specific filtering mediated by the hear/torso/pinnae, using the head-related transfer functions from the standard KEMAR mannequin. Neural networks were trained to classify the location of the sound source (azimuth and elevation) into one of a large set of small bins (spaced 5 degrees in azimuth and 10 degrees in elevation). C. Example neural network architectures from architecture search. Architectures consisted of sequences of “blocks” (a normalization layer, followed by a convolution layer, followed by a nonlinearity layer) followed by a pooling layer, culminating in fully connected layers followed by a classifier that provided the network’s output. Architectures varied in the total number of layers, the kernel dimensions for each convolutional layer, the number of blocks that preceded each pooling layer, and the number of fully connected layers preceding the classifier. Labels indicate an example block, pooling layer, and fully connected layer.

In the absence of models that can instantiate localization behavior, the science of sound localization has largely relied on intuitions about optimality. Those intuitions were invaluable in stimulating research, but in some cases proved incorrect. For instance, it was supposed for many decades that interaural cues neatly segregated in their function, with time and level differences subserving localization for low- and high-frequency sounds, respectively (“duplex” theory) (14). However, it has become clear that level differences are perceptually important even at low frequencies (35–37). Here and in other domains of sensory processing, models that can replicate real-world behavior, and thus help reveal the information needed to mediate such behavior, could have a transformative effect on perceptual science.

Here we exploit the power of contemporary artificial neural networks to develop a model optimized to localize sounds in realistic conditions. Unlike much other contemporary work using neural networks to investigate perceptual systems (38–43), our primary interest is not in potential correspondence between internal representations of the network and the brain. Instead, we aim to use the neural network as a way to find an optimized solution to a difficult real-world task that is not easily specified analytically. Our approach is thus analogous to the classic ideal observer approach, but harnesses modern machine learning to approximate an ideal observer for a problem where one is not analytically tractable.

To give the network access to the same cues available to biological organisms, we trained it on a high-fidelity cochlear representation of sound, leveraging recent technical advances (44) to train the large models that are required for such high-dimensional input. Unlike previous generations of neural network models (21), which were reliant on hand-specified sound features, we learn all subsequent stages of a sound localization system to obtain good performance in real-world conditions. To obtain sufficent labeled data with which to train the network, we used a virtual acoustic world (45) to simulate sounds at different locations with realistic patterns of surface reflections and background noise.

The resulting model replicated a large and diverse array of human behavioral characteristics. We then performed experiments on the model that are impossible with biological organisms, training it in unnatural conditions to simulate evolution and development in alternative worlds. These experiments suggest that the characteristics of human hearing are indeed adapted to the constraints of real-world localization, and that the rich panoply of sound localization phenemona can be explained as consequences of this adaptation. The approach we employ is broadly applicable to other sensory modalities and problems.

## Results

### Model Construction

We began by building a system that could localize sounds using the information available to human listeners. The system thus had outer ears (pinnae), and a simulated head and torso, along with a simulated cochlea. The outer ears and head/torso were simulated using head-related impulse responses recorded from a standard physical model of the human (46). The cochlea was simulated with a bank of bandpass filters modeled on the frequency selectivity of the human ear (47), whose output was rectified and downsampled to 4 kHz to simulate the upper limit of phase locking in the auditory nerve (48). The inclusion of a fixed cochear front-end (in lieu of trainable filters) reflected the assumption that the cochlea evolved to serve many different auditory tasks rather than being primarily driven by sound localization. As such, the cochlea seemed a plausible biological constraint on sound localization.

The output of the two cochlea formed the input to a convolutional neural network (Figure 1B). This network instantiated a cascade of simple operations – filtering, pooling, and normalization, the parameters of which were tuned to maximize performance on the training task. The optimization procedure had two nested phases: an architecture search in which we searched over architectural parameters to find a network architecture that performed well (Figure 1C), and a training phase in which the filter weights of the selected architectures were trained to asymptotic performance levels using gradient descent.

The architecture search consisted of training each of a large set of possible architectures for 15000 training steps with 16 stimulus examples per step (240k total examples; see Supplemental Figure 1 for distribution of localization performance across architectures). We then chose the 10 networks that perfomed best on a validation set of data not used during training (Supplemental Table 1). The parameters of these 10 networks were then reinitialized and each trained for 100k training steps (1.6M examples). Given evidence that internal representations can vary somewhat across different networks trained on the same task (49), we present results aggregated across the top 10 best-performing architectures, treated akin to different participants in an experiment. Most results graphs present the average results for these 10 networks, which we collectively refer to as “the model”.

The training data was based on a set of ~500,000 stereo audio signals with associated 3D locations relative to the head. These signals were generated from 385 2-second natural sound source recordings rendered at a spatial location in a simulated room. The room simulator used a modified source-image method (45, 50) to simulate the reflections off the walls of the room. These reflections were then filtered by the head-related impulse response for the direction of the reflection (46). Five different rooms were used, varying in their dimensions and in the material of the walls. To mimick the common presence of noise in real-world environments, each training signal also contained spatialized noise. Background noise was synthesized from the statistics of a natural sound texture (51), and was rendered at several randomly chosen locations using the same room simulator. At each training step the rendered natural sound sources were randomly paired with rendered background noises.The neural networks were trained to map the binaural audio to the location of the sound source (specified by its azimuth and elevation relative to the network’s “head”).

### Model Evaluation in Real-World Conditions

The trained networks were first evaluated on a held-out set of 70 sound sources rendered using the same pipeline used to generate the training data (yielding a total of ~47,000 stereo audio signals). The best-performing networks produced accurate localization for this validation set (the mean error was 5.3 degrees in elevation and 4.4 degrees in azimuth, front-back folded, i.e. reflected about the coronal plane, to discount front-back confusions).

To assess whether the networks would generalize to real-world stimuli outside the training distribution, we made binaural recordings in an actual conference room using a mannequin with in-ear microphones (Fig. 2A). Humans localize relatively well in such free-field conditions (Figure 2B). The trained networks also localized real-world recordings relatively well (Figure 2C), on par with human free-field localization, with errors limited to the front-back confusions that are common to humans when they cannot move their heads (Figure 2D) (52, 53).

**Figure 2.**
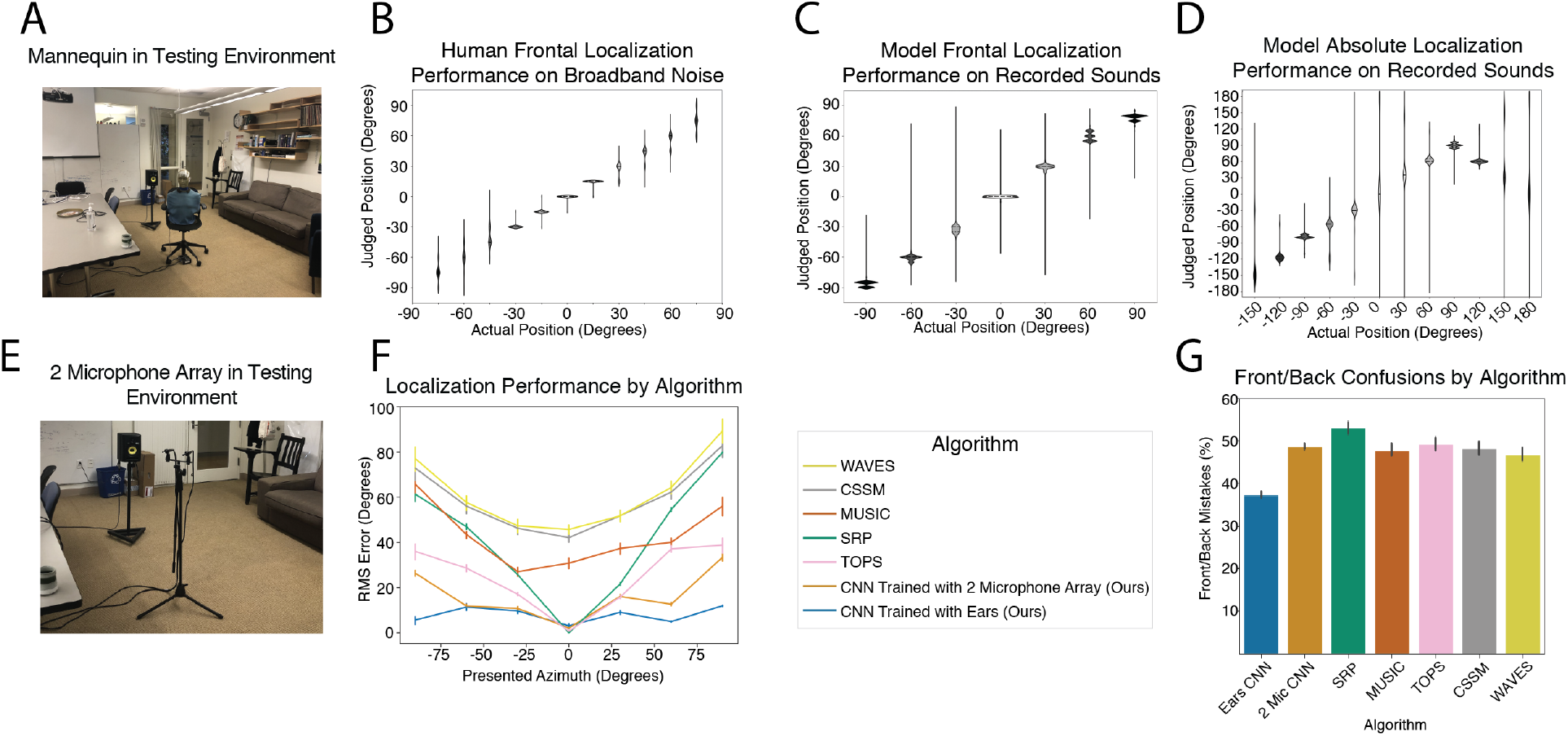
Model performance in real-world environments. A. Photo of recording setup for real-world test set. The mannequin was seated on a chair that was rotated relative to the speaker to achieve different azimuthal positions. Sound was recorded from microphones in the mannequin ears. B. Free-field localization of human listeners. Participants heard a sound played from one of 11 speakers in the front horizontal plane and pointed to the location. Data are replotted from a previous publication (54). C. Localization judgments of the neural networks trained with simulated ears/head/torso. The input to the networks were recordings from a mannequin’s ears made in the conference room shown in A. Graph plots kernel density estimates of response distribution. For ease of comparison with human experiment in B, in which all locations were in front of the listener, positions were front-back folded (both actual and judged positions were reflected across the coronal plane). D. Localization judgments of the neural networks without front-back folding. Network errors are predominantly at front-back reflections of the correct location. E. Photo of two-microphone array. Microphone spacing was the same as in the mannequin, but the recordings lacked the acoustic effects of the pinnae, head, and torso. F. Localization accuracy of standard two-microphone localization algorithms, our neural network localization model trained with ear/head/torso filtering effects (same as 2B and 2D), neural networks trained instead with simulated input from the two-microphone array, and human listeners (calculated from the data in 2B, obtained in free-field anechoic noise-free conditions). Localization judgments are front-back folded. Error bars here and in G plot SEM, obtained by bootstrapping across stimuli. G. Front-back confusions by each of the algorithms from F. Chance level is 50%.

For comparison, we also assessed the performance of a standard set of two-microphone localization algorithms from the engineering literature (55–59). In addition, we trained a neural network to localize sounds from a two-microphone array that lacked the filtering provided to biological organisms by the ears/head/torso but that included the simulated cochlea (Figure 2E). Our networks that had been trained with biological pinnae/head/torso outperformed the set of standard two-microphone algorithms from the engineering community, as well as the neural network trained with stereo microphone input without a head and ears (Figure 2F&G). This latter result confirms that the head and ears provide valuable cues for localization. Overall, performance on the real-world test set demonstrates that training a neural network in a virtual world produces a model that can accurately localize sounds in realistic conditions (to our knowledge the first such model in existence).

### Model Behavioral Characteristics

To assess whether the trained networks replicated the characteristics of human sound localization, we simulated a large set of experiments from the literature, intended to span the best-known and largest effects in spatial hearing. We emphasize that the networks were not fit to human data in any way. Despite this, the networks reproduced the characteristics of human spatial hearing across this broad set of experiments.

#### Sensitivity to interaural time and level differences

We began by assessing whether the networks learned to use the binaural cues known to be important for biological sound localization. We probed the effect of interaural time and level differences (ITDs and ILDs, respectively) on localization behavior using a paradigm in which additional time and level differences are added to sounds with different frequency content rendered in virtual acoustic space (60) (Figure 3A, left). This paradigm has the advantage of using realistically externalized sounds and an absolute localization judgment (rather than the left/right lateralization judgments of more simpler stimuli that are common to many other experiments (61–64)).

**Figure 3.**
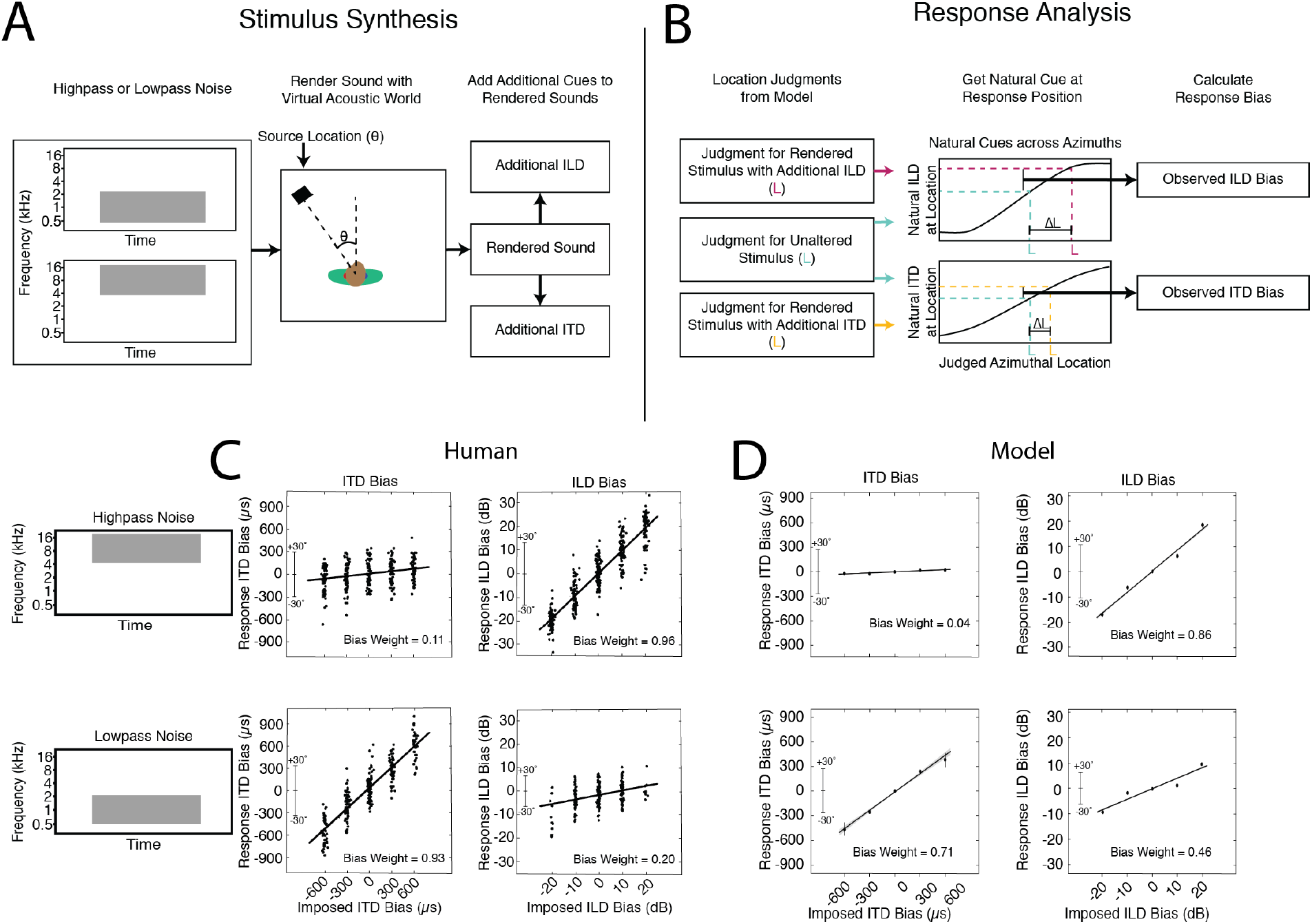
Sensitivity to interaural time and level differences. A. Schematic of stimulus generation. Noise bursts filtered into high or low frequency bands were rendered at a particular azimuthal position, after which an additional ITD or ILD was added to the stereo audio signal. B. Schematic of response analysis. Responses were analyzed to determine the amount by which the perceived location was altered (ΔL) by the additional ITD/ILD, expressed as the amount by which the ITD/ILD would have changed if the actual sound’s location changed by ΔL. C. Effect of additional ITD and ILD on human localization. Y axis plots amount which the perceived location is altered expressed in ITD/ILD, as described above. Data reproduced from a previous publication (60) with permission of the authors. D. Effect of additional ITD and ILD on model localization. Same conventions as B. Error bars plot SEM, bootstrapped across the 10 networks.

In the original experiment (60), the change to perceived location imparted by the additional ITD or ILD was expressed as the amount by which the ITD or ILD would change in natural conditions if the actual location were changed by the perceived amount (Figure 3A, right). This yields a curve whose slope indicates the efficacy of the manipulated cue (ITD or ILD). We reproduced the stimuli from the original study, rendered them in our virtual acoustic world, added ITDs and ILDs as in the original study, and analyzed the model’s localization judgments in the same way.

For human listeners, ITD and ILD have opposite efficacies at high and low frequencies (Figure 3B), as predicted by classical duplex theory. An ITD bias imposed on low-frequency sounds shifts the perceived location of the sound substantially (bottom left), whereas an ITD imposed on high-frequency sound does not (top left). The opposite effect occurs for ILDs (right panels), although there is a weak effect of ILDs on low-frequency sound. This latter effect is inconsistent with the classical duplex story (14) but consistent with more modern measurements indicating small but reliable ILDs at low frequencies (37) that are used by the human auditory system (35, 36, 65).

As shown in Figure 3C, the model results qualitatively replicate the effects seen in humans. Added ITDs and ILDs have the largest effect at low and high frequencies, respectively, but ILDs have a modest effect at low frequencies as well. This produced an interaction between the type of cue (ITD/ILD) and frequency range (difference of differences between slopes significantly greater than 0; p<.00001, evaluated by bootstrapping across the 10 networks). However, the effect of ILD at low frequencies was also significant (slope significantly greater than 0; p<.00001, via bootstrap). Thus, a model optimized for accurate localization both exhibits the dissociation classically associated with duplex theory, but also its refinements in the modern era.

#### Azimuthal localization of broadband sounds

We next measured localization accuracy of broadband noise rendered at different azimuthal locations (Figure 4A). In humans, localization is most accurate near the midline (Figure 4B), and becomes progressively less accurate as sound sources move to the left or right of the listener (66–68). One explanation is that the first derivatives of ITD and ILD with respect to azimuthal location decrease as the source moves away from the midline (18), providing less information about location (25). The network qualitatively reproduces this result (Figure 4C).

**Figure 4.**
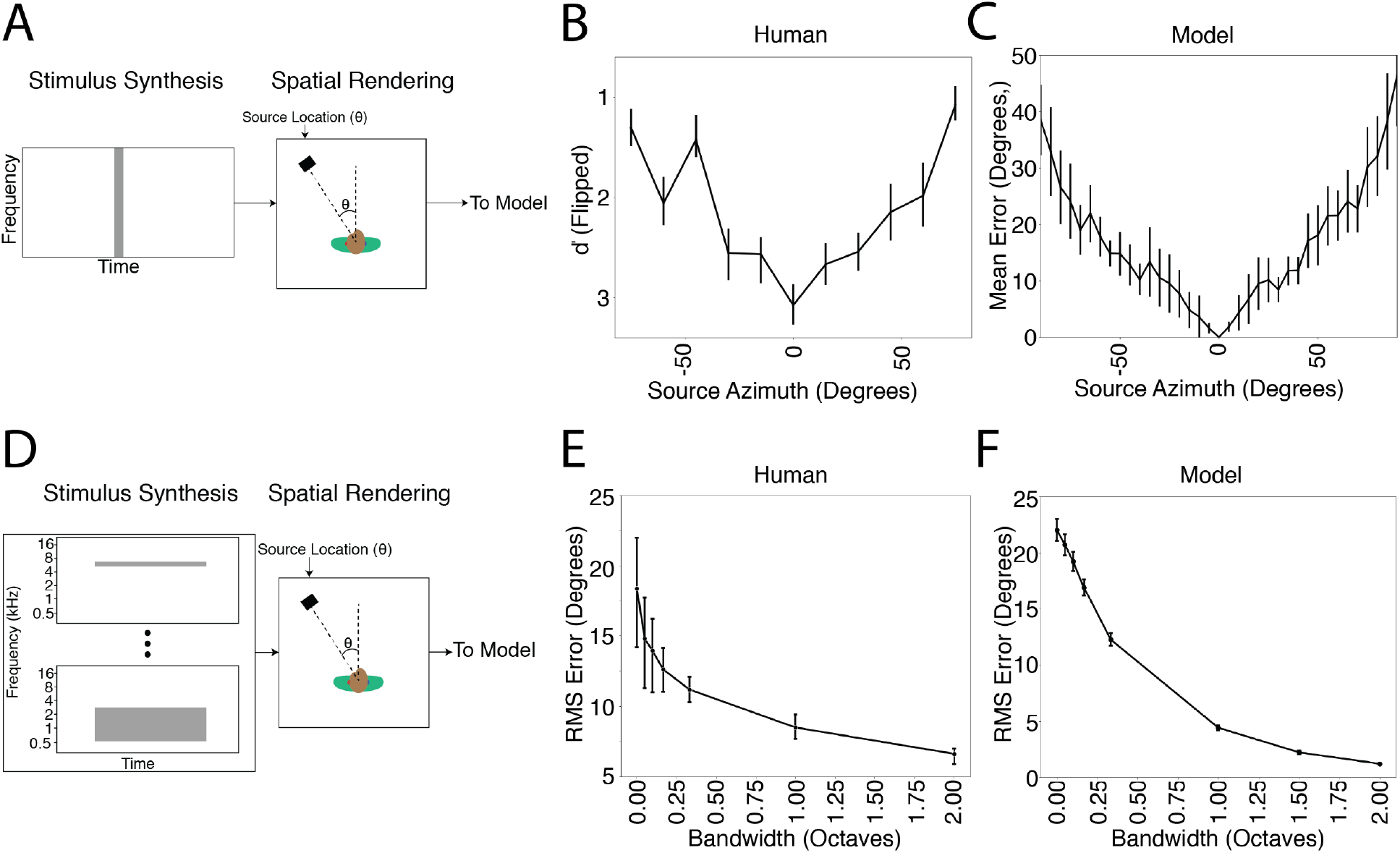
Azimuthal localization is most accurate at the midline and improves with stimulus bandwidth. A. Schematic of stimuli and task measuring localization accuracy at different azimuthal positions. B. Localization accuracy of human listeners for broadband noise at different azimuthal positions. Data were scanned from a previous publication (68). C. Model accuracy for broadband noise at different azimuthal positions. Error bars plot SEM across the 10 networks. D. Schematic of stimuli and task measuring effect of bandwidth on localization accuracy. Noise bursts varying in bandwidth were presented at particular azimuthal locations; participants indicated the azimuthal position with a keypress. E. Effect of bandwidth on human localization of noise bursts. Data are replotted from a previous publication (69). F. Effect of bandwidth on model localization of noise bursts. Networks were constrained to report only the azimuth of the stimulus. Error bars plot SEM across the 10 networks.

#### Integration across frequency

Because biological hearing begins with a decomposition of sound into frequency channels, binaural cues are believed to be initially extracted within these channels (17, 22). However, organisms are believed to integrate information across frequency to achieve more accurate localization than could be mediated by any single frequency channel. One signature of this integration is improvement in localization accuracy as the bandwidth of a broadband noise source is increased (Figure 4D&E) (69, 70). We replicated one such experiment on the networks and they exhibited a similar effect, with accuracy increasing with noise bandwidth (Figure 4F).

#### Use of ear-specific cues to elevation

In addition to the binaural cues that provide information about azimuth, organisms are known to make use of the direction-specific filtering imposed on sound by the ears, head and torso (15, 71). Each individual’s ears have resonances that “color” a sound differently depending on where it comes from in space. Individuals are believed to learn the specific cues provided by their ears. In particular, if forced to listen with altered ears, either via molds inserted into the ears (72) or via recordings made in a different person’s ears (73), localization in elevation degrades even though azimuthal localization is largely unaffected (Figure 5A-C).

**Figure 5.**
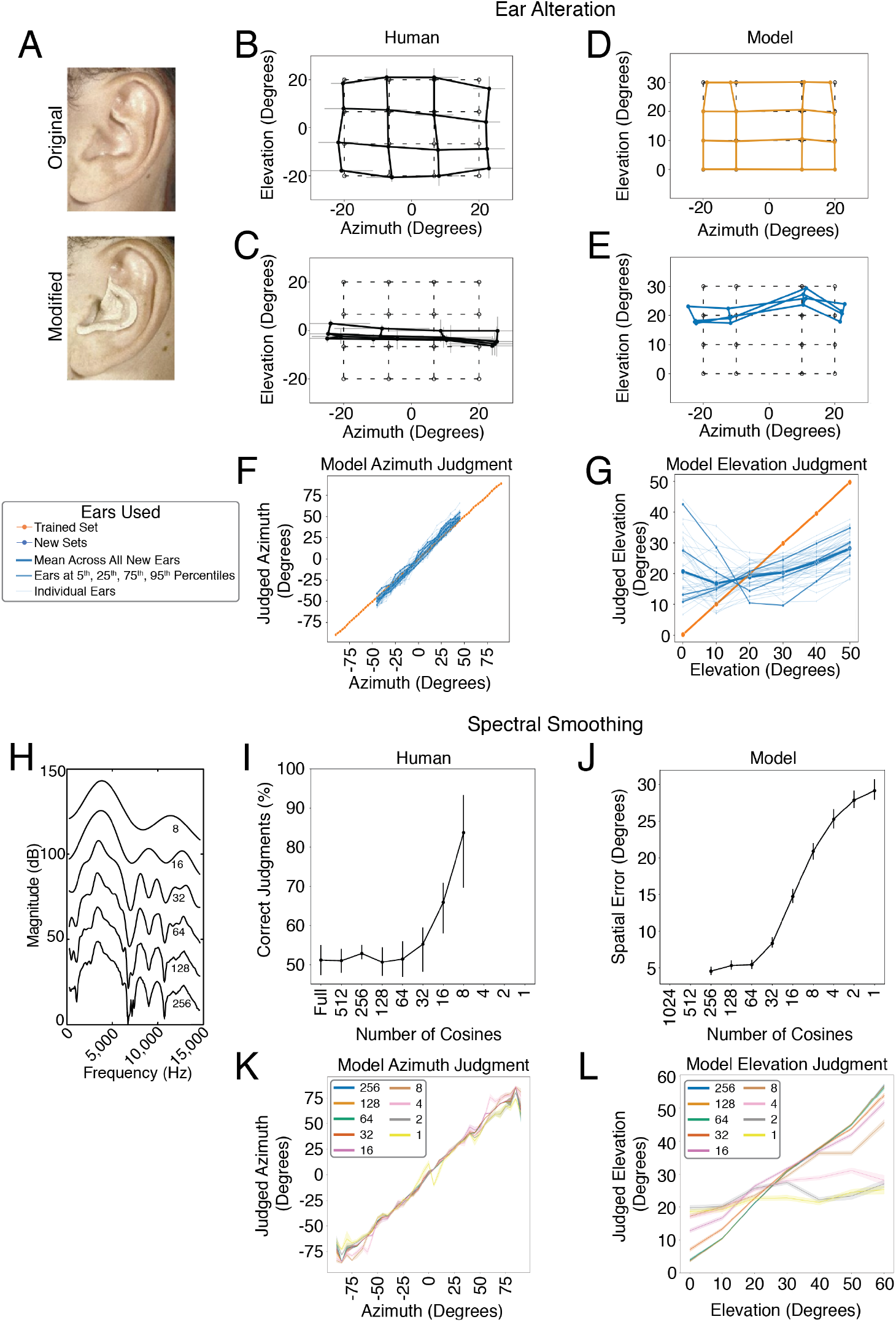
Dependence of elevation perception on ear-specific transfer functions. A. Photographs of example ear alteration in humans (reproduced from a previous publication (72) with permission of the authors). B. Sound localization in azimuth and elevation by human listeners with unmodified ears, who saccaded to the perceived location of a broadband white noise source in a dark anechoic room. Figure plots mean and SEM of perceived locations for 4 participants, superimposed on grid of true locations (dashed lines). Data scanned from original publication (72). C. Effect of ear alteration on human sound localization. Same conventions as B. D. Sound localization in azimuth and elevation by the model, using the ears (head-related impulse responses) from training, with broadband white noise sound sources. Tested locations differed from those in the human experiment in order to conform to the location bins used for network training. E. Effect of ear alteration on model sound localization. Ear alteration was simulated by substituting an alternative set of head-related impulse responses into the sound rendered following training. Graph plots average results across all 45 sets of alternative ears. F. Effect of individual sets of alternative ears on localization in azimuth. Graph shows results for a larger set of locations than in D and E to illustrate the generality of the effect. Each thin blue line shows results for a different set of ears (head-related impulse responses). Bolded lines show ears at 5^th^, 25^th^, 75^th^, and 95^th^ percentiles when the 45 sets of ears were ranked according to accuracy. Thick bold line shows mean across all 45 sets. G. Effect of individual sets of alternative ears on localization in elevation. Same conventions as in F. H. Smoothing of head-related transfer functions, produced by varying the number of coefficients in a discrete cosine transform of the transfer function. Reproduced from original publication (74) with permission of the authors. I. Effect of spectral smoothing of head-related transfer functions on human perception. Human listeners heard two sounds, one played from a speaker in front of them, and one played through open-backed earphones, and judged which was played through earphones. The earphone-presented sound was rendered using HRTFs smoothed by various degrees. In practice participants performed the task by noting changes in sound location. Data scanned from original publication (74). Conditions with 4, 2, and 1 cosine components were omitted from the experiment, but are included on the x axis to facilitate comparison with the model results in J. J. Effect of spectral smoothing on model sound localization accuracy (measured in both azimuth and elevation). Conditions with 512 and 1024 cosine components were not realizable given the length of the impulse responses we used, but are included on the x axis to facilitate comparison with the human results in I. K. Effect of spectral smoothing on model accuracy evaluated separately in azimuth. L. Effect of spectral smoothing on model accuracy evaluated separately in elevation.

To test whether the trained networks similarly learned to use ear-specific elevation cues, we measured localization accuracy in elevation and azimuth. We compared two conditions: one where sounds were rendered using the head-related impulse response set used for training the networks, and another where the impulse responses were different (having been recorded in a different person’s ears). Because we have unlimited ability to run experiments on the networks, in the latter case we evaluated localization with 45 different sets of impulse responses, each recorded from a different human. As expected, localization of sounds rendered with the ears used for training was good in both azimuth and elevation (Figure 5D). But when tested with different ears, localization in elevation generally collapsed (Figure 5E), much like what happens to human listeners when molds are inserted in their ears (Figure 5C), even though azimuthal localization was nearly indistinguishable from that with the trained ears. Results for individual sets of alternative ears reveal that elevation performance transfers better across some ears than others (Figure 5F), consistent with anecdotal evidence that sounds rendered with HRTFs other than one’s own can sometimes be convincingly localized in three dimensions.

#### Limited spectral resolution of elevation cues

Elevation perception is believed to rely on the peaks and troughs introduced to a sound’s spectrum by the ears/head/torso (Figure 1A, right). In humans, however, perception is dependent on relatively coarse spectral features – the transfer function can be smoothed substantially before human listeners notice abnormalities (74) (Figure 5G & H), for reasons that are unclear. In the original experiment, human listeners discriminated sounds with and without smoothing, a judgment that was in practice made by noticing changes in the apparent location of the sound. To test whether the trained networks exhibited a similar effect, we presented sounds to the networks with similarly smoothed transfer functions and measured the extent to which the localization accuracy was affected. The effect of spectral smoothing on the networks’ accuracy is similar to the measured sensitivity of human listeners (Figure 5I). The effect of the smoothing is most prominent for localization in elevation, as expected, but there is also some effect on localization in azimuth for the more extreme degrees of smoothing (Figure 5J), consistent with evidence that spectral cues affect azimuthal space encoding (75).

#### The precedence effect

Another hallmark of biological sound localization is that judgments are biased towards information provided by sound onsets (18, 76). The classic example of this bias is known as the “precedence effect” (77–79). If two clicks are played from speakers at different locations with a short delay (Figure 6A), listeners perceive a single sound whose location is determined by the click that comes first. The effect is often hypothesized to be an adaptation to the common presence of reflections in real-world listening conditions – reflections traverse longer paths and thus arrive later, and from an erroneous direction (18). To test whether our model would exhibit a similar effect, we simulated the classic precedence experiment, rendering two clicks at different locations. When presented simultaneously, the model reports the sound to be centered between the two click locations, but when a small delay is introduced, the reported location is that of the leading click (Figure 6B). This effect breaks down as the delay is increased, as in humans, though with the difference that the network cannot report hearing two sounds, and so instead reports a single location is intermediate between those of the two clicks.

**Figure 6.**
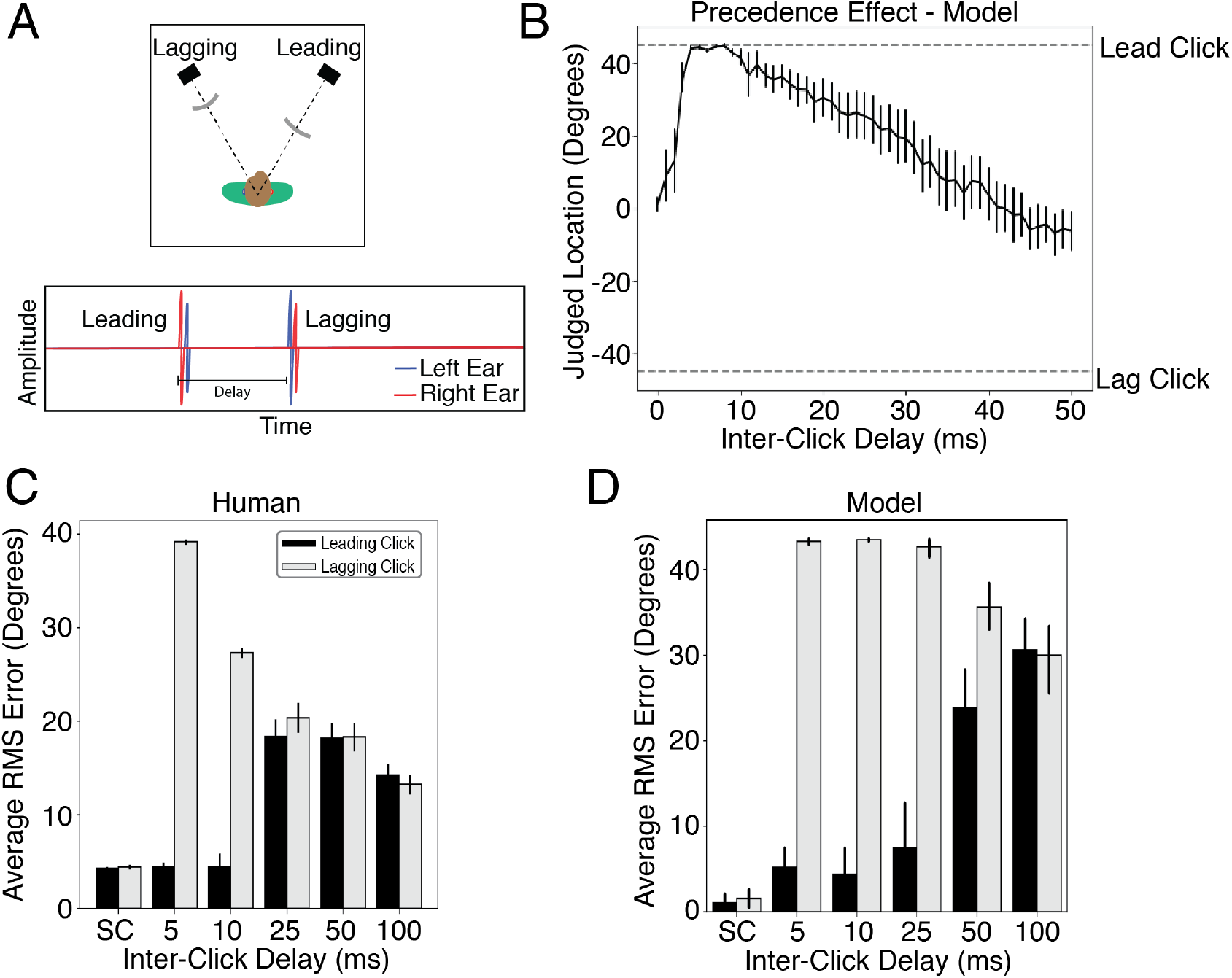
The precedence effect. A. Diagram of stimulus. Two clicks are played from two different locations relative to the listener. The time interval between the clicks is manipulated and the listener is asked to localize the sound(s) that they hear. When the delay is short but non-zero, listeners perceive a single click at the location of the first click. At longer delays listeners hear two distinct sounds. B. Localization judgments of the model for two clicks at +45 and −45 degrees. The model exhibits a bias for the leading click when the delay is short but non-zero. At longer delays the model judgments (which are constrained to report the location of a single sound, unlike humans), converge to the average of the two click locations. Error bars plots SEM across the 10 networks. C. Error in localization of the leading and lagging clicks by humans as a function of delay. SC denotes a single click at the leading or lagging location. Data scanned from original publication (80). D. Error in localization of the leading and lagging clicks by the model as a function of delay. Error bars plots SEM across the 10 networks.

To compare the model results to human data, we simulated an experiment in which participants reported the location of both the leading and lagging click as the inter-click delay was varied (80). At short but non-zero delays, humans accurately localize the leading but not the lagging click (Figure 6C; because a single sound is heard at the location of the leading click). At longer delays the lagging click is more accurately localized, and listeners start to mislocalize the leading click, presumably because they confuse which click is first and so mis-match their responses (80). The model qualitatively replicates both effects, in particular the large asymmetry in localization accuracy for the leading and lagging sound at short delays (Figure 6D).

#### Summary of psychophysical results

Despite having no previous exposure to the tone and noise stimuli used in the experiments, the model qualitatively replicated a wide range of classic behavioral effects found in humans. We emphasize that the networks were not fit to match human data in any way. They were optimized only for accurate localization of natural sounds in realistic conditions – an entirely different stimulus set from that used in the simulated experiments shown in Figures 3-6. The human-like sound localization behaviors in the model emerge as side effects of this optimization. These results raise the possibility that the characteristics of biological sound localization may be understood as a consequence of optimization for real-world localization. However, given these results alone, the role of the natural environment in determining these characteristics is left unclear.

### Effect of Optimization for Unnatural Environments

To assess the extent to which the properties of biological spatial hearing are adapted to the constraints of localization in natural environments, we took advantage of the ability to use our network training procedure to conduct experiments that are impossible with biological organisms, training the networks in virtual worlds altered in various ways to simulate the effects of evolution and/or development in alternative environments (Figure 7A). We altered the training environment in one of three ways: 1) by eliminating reflections (simulating surfaces that absorb all sound that reaches them, unlike real-world surfaces), 2) by eliminating background noise, and 3) by replacing natural sound sources with artificial sounds (narrowband noise bursts). In each case we trained the networks to asymptotic performance, and then ran them on the full suite of psychophysical experiments described above. We then quantified the similarity between the model psychophysical results and those of humans as the mean squared error between the model and human results on each experiment (see Methods).

**Figure 7.**
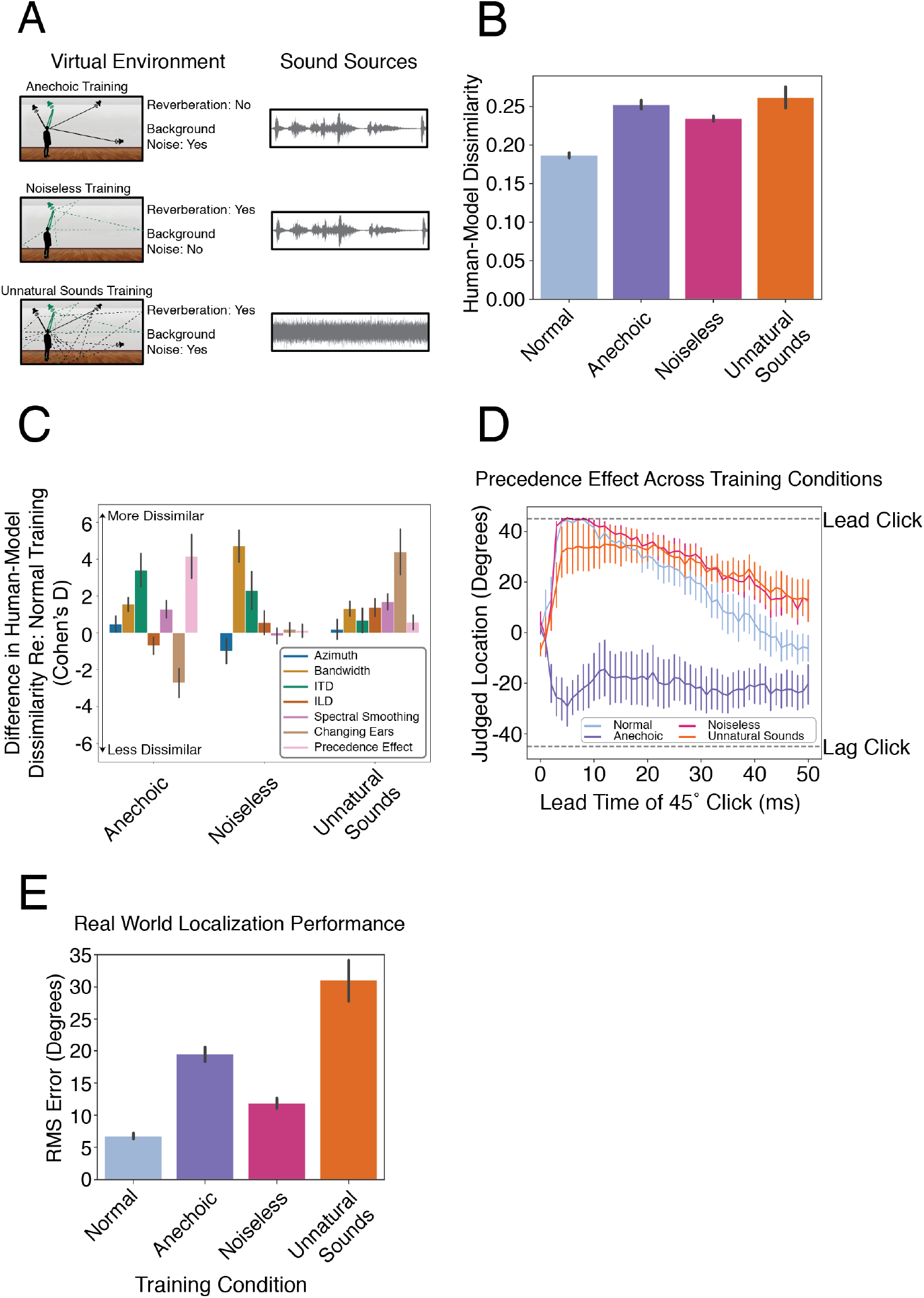
Effect of unnatural training conditions. A. Schematic depiction of altered training conditions, eliminating echoes or background noise, or using unnatural sounds. B. Overall human-model dissimilarity for natural and unnatural training conditions. Error bars plot SEM, bootstrapped across networks. C. Effect of unnatural training conditions on human-model dissimilarity for individual experiments, expressed as the effect size of the difference in dissimilarity between the natural and each unnatural training condition (Cohen’s D, computed between human-model dissimilarity for networks in normal and modified training conditions). Positive numbers denote a worse resemblance to human data compared to the model trained in normal conditions. Error bars plot SEM, bootstrapped across the 10 networks D. The precedence effect in networks trained in alternative environments. E. Real-world localization accuracy of networks for each training condition. Error bars plot SEM, bootstrapped across the 10 networks.

Figure 7B shows the average dissimilarity between the human and model results on the suite of psychophysical experiments, computed separatedly for each model training condition. The dissimilarity is lowest for the model trained in natural conditions, and significantly higher for each of the alternative conditions (p<.00001 in each case, via bootstrap across the 10 networks; results were fairly consistent across networks, Supplemental Figure 4). This result provides additional evidence that the properties of spatial hearing are consequences of adaptation to the natural environment – human-like spatial hearing emerged from task optimization only for naturalistic training conditions.

To get insight into how the environment influences perception, we examined the human-model dissimilarity for each experiment individually (Figure 7C). Because the absolute dissimilarity is not meaningful (in that it is limited by the reliability of the human results, which is not perfect), we assessed the differences in human-model dissimilarity between the natural training condition and each unnatural training condition. These differences were most pronounced for a subset of experiments in each case. The anechoic training condition produced most abnormal results for the precedence effect, but also produced substantially different results for ITD cue strength (the effect size for the difference in human-model error between anechoic and natural training conditions was significantly greater in both these two experiments than in the other experiments, p<0.0001, via bootstrap across networks). The noiseless training condition produced most abnormal results for the effect of bandwidth (p<0.0001, via bootstrap across networks). The training condition with unnatural sounds produced most abnormal results for the experiment measuring the effects of alternative ears (p<0.0001, via bootstrap across networks), presumably because without the pressure to localize broadband sounds, the model did not acquire sensitivity to spectral cues to elevation. These results indicate that different worlds would lead to different perceptual systems with distinct localization strategies.

The most interpretable example of environment-driven localization strategies is the precedence effect. This effect is shown in Figure 7D for models trained in each of the four virtual environments. Anechoic training completely eliminates the effect, even though it is largely unaffected by the other two unnatural training conditions, substantiating the hypothesis that the precedence effect is an adaptation to reflections in real-world listening conditions. See Supplementary Figures 2 and 3 for full psychophysical results for models trained in alternative conditions.

In addition to diverging from the perceptual strategies found in human listeners, the networks trained in unnatural conditions performed more poorly at real-world localization. When we ran networks trained in alternative conditions on our real-world test set of recordings from mannequin ears in a conference room, localization accuracy was substantially worse in all cases (Figure 7E; p<.0.0001 in all cases). Coupled with the abnormal psychophysical results of these alternatively trained networks, this result indicates that the classic perceptual characteristics of spatial hearing reflect strategies that are important for real-world localization, in that systems that deviate from these characteristics localize poorly.

### Model Predictions of Sound Localizability

One advantage of a model that can mediate actual localization behavior is that one can run large numbers of experiments on the model, searching for “interesting” predictions that might then be tested in human listeners. Here we used the model to estimate the accuracy with which different natural sounds would be localized in realistic conditions. We chose to examine musical instrument sounds as these are both diverse and available as clean recordings in large numbers. We took a large set of instrument sounds (81) and rendered them at a large set of randomly selected locations. We then measured the average localization error for each instrument.

As shown in Figure 8A, there was reliable variation in the accuracy with which instrument sounds were localized by the model. The median error was as low as 1.06 degrees for Reed Instrument #3 and as high as 40.02 degrees for Mallet #1, folded to discount front-back confusions (without front-back folding the overall error was larger, but the ordinal relations among instruments was similar). The human voice was also among the most acurately localized sounds in the set we examined, with a mean error or 2.39 degrees (front-back folded).

**Figure 8.**
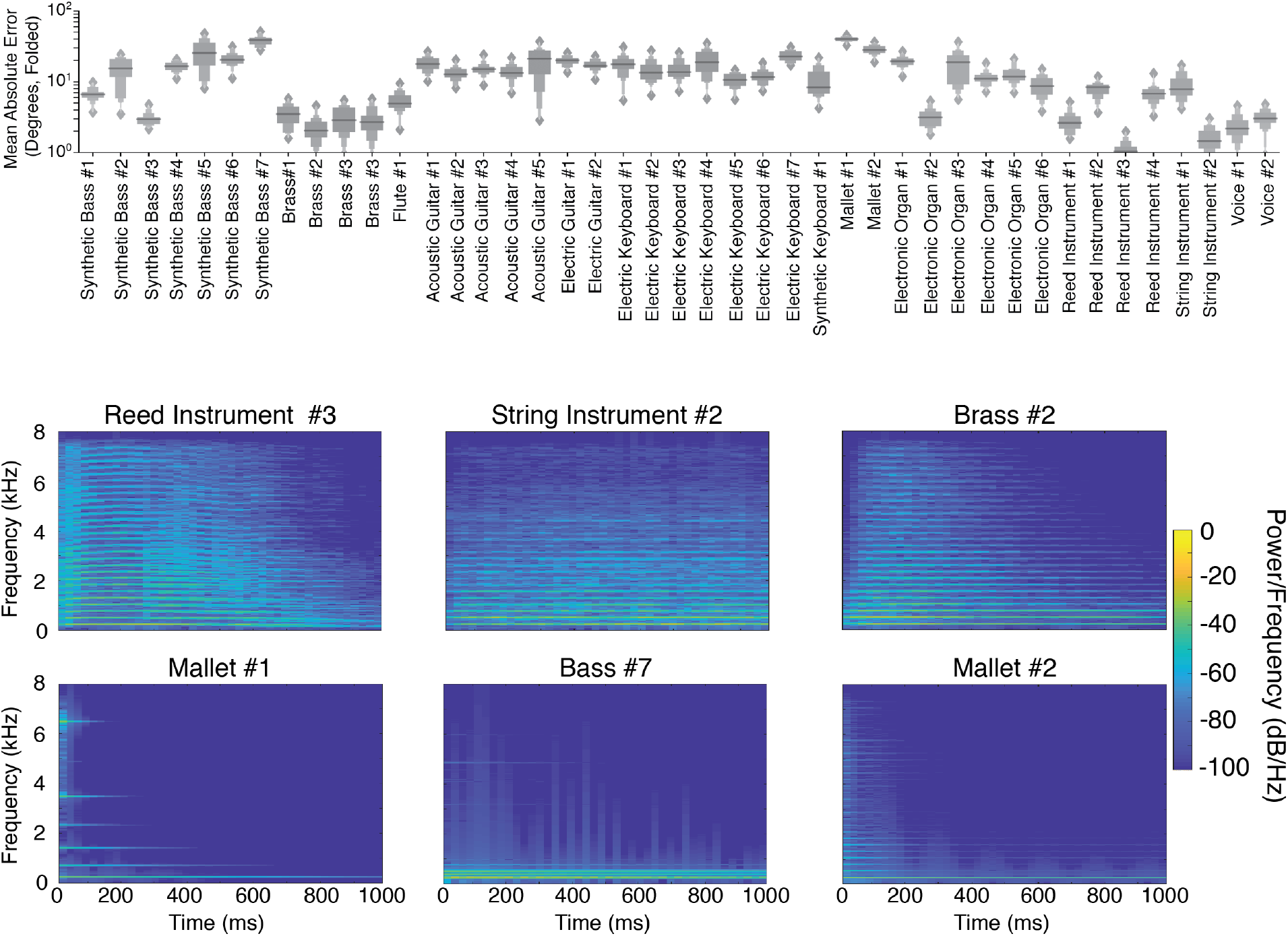
Model localization accuracy for musical instrument sounds. A. Mean model localization error for each of 43 musical instruments. Each of a set of instrument notes was rendered at randomly selection locations. Graph shows box plots of the mean localization error across notes, measured after actual and judged positions were front-back folded. B. Spectrograms of example notes (middle C) for the three most and least accurately localized instruments (top and bottom, respectively).

Figure 8B displays spectrograms for example notes for the three best- and worst-localized instruments. It is apparent that the best-localized instruments are spectrally dense, and thus presumably take advantage of cross-frequency integration and ear-specific spectral cues (which improve localization accuracy in both humans and the model; Figure 4E&F, and Figure 6). This result is consistent with the common idea that narrowband sounds are less well localized, but the model provides a quantitative metric of localizability that we would not otherwise have.

To assess whether the results could be predicted by simple measures of spectral sparsity, we measured the spectral flatness of each instrument sound (the ratio of the geometric mean of the power spectrum to the arithmetic mean of the power spectrum). The average spectral flatness of an instrument was significantly correlated with the model’s localization accuracy (r_s_ = .77, p<.0001), but this correlation was well below the split-half reliability of the model’s accuracy for an instrument (r_s_ = .99). This difference suggests that there are sound features above and beyond spectral density that determine a sound’s localizability, and illustrates the value of an optimized system to make perceptual predictions.

We had intentions of running a free-field localization experiment in humans to test these predictions, but had to halt experiments due to COVID-19. We have hopes of running the experiment in the future. However, we note that informal observation indicates that the model predictions hold true to some extent. The first author tested himself by spinning himself around in a rotating chair while blindfolded, and trying to point to a speaker playing an instrument note when the chair came to a halt. The sounds that are poorly localized by the model were also quite difficult for the first author to localize.

## Discussion

We trained artificial neural networks to localize sounds from binaural audio rendered in a virtual world and heard through simulated ears. When the virtual world mimicked natural auditory environments, with surface reflections, background noise, and natural sound sources, the trained networks replicated many attributes of spatial hearing found in biological organisms. These included the frequency-dependent use of interaural time and level differences, the use of ear-specific spectral cues to elevation and robustness to spectral smoothing of these cues, the integration of spatial information across frequency, and localization dominance of sound onsets. The model successfully localized sounds in an actual real-world environment better than alternative algorithms that lacked ears. The model also made predictions about the accuracy with which different types of real-world sounds could be localized. But when the training conditions were altered to deviate from the natural environment by eliminating surface reflections, background noise, or natural sound source structure, the behavioral characteristics of the model deviated notably from human-like behavior. The results suggest that most of the key properties of mammalian spatial hearing can be understood as consequences of optimization for the task of localizing sounds in natural environments. Our approach extends classical ideal observer analysis to new domains, where provably optimal analytic solutions are difficult to attain but where supervised machine learning can nonetheless provide optimized solutions in different conditions.

### Contributions Relative to Previous Work

We provide the first model of biological sound localization that can actually localize sounds in real-world settings. The ability to localize in realistic conditions was enabled by learning a solution to the localization problem from data. Prior models of sound localization required cues to be hand-coded and provided to the model by the experimenter (19–21, 31). In some cases previous models were able to derive optimal encoding strategies for such cues (25), which could be usefully compared to neural data (82). In other cases models were able to make predictions of behavior in simplified conditions using idealized cues (31). However, the idealized cues that such models work with are not well-defined for arbitrary real-world stimuli, preventing the modeling of general localization behavior. It has thus not previously been possible to derive optimal behavioral characteristics for real-world behavior.

Our results highlight the power of contemporary machine learning to achieve realistic behavioral competence in computational models. The fact that our model can mediate realistic localization behavior greatly expands the scope of what it can be used for. In particular, the model is able to perform any experiment that relies on localization judgments, and can make predictions about real-world behavior with natural sounds. The model’s real-world behavioral competence also enabled a direct demonstration of the utility of having ears on a head – alternative systems equipped with two microphones performed substantially worse.

Another opportunity afforded by deriving perceptual strategies from data is that one can examine the constraints that yield biological solutions (83, 84). We found that when the model was optimized for unnatural conditions it yielded alternative solutions with distinct behavioral signatures (Figure 7). The results suggest that many aspects of behavior can be explained by task and environmental constraints rather than algorithmic or mechanistic characteristics of the underlying neural processes. Learned task solutions thus provide a way to link perception to the environment in situations that may be practically impossible to test with humans. The approach provides an additional tool with which to understand evolution (85), and a way to link experimental results with function.

Our model provides normative explanations for a broad suite of spatial hearing characteristics. In some cases these had been hypothesized but not definitively established, as with the precedence effect, which our results suggest is indeed an adaptation to reverberation (Figure 7D). In other cases the model provided explanations for behavioral characteristics that previously had none, such as the relatively coarse spectral resolution of elevation perception (Figure 5I&J), which evidently reflects the absence of reliable information at finer resolutions.

To our knowledge our model is the first example of a model trained entirely in a virtual environment that transfers to real-world conditions. Although contemporary supervised learning has been embraced by the computational neuroscience community as a way to generate models of brain function, one of its major limitations is the need for large amounts of labeled data, typically acquired via painstaking human annotation. Virtual environments allow the scientist to generate the data, with the labels coming for free (as the parameters used to generate the data). Virtual training environments have the potential to greatly expand the settings in which supervised learning can be used to develop models of the brain. The approach has been increasingly adopted in the engineering world (86, 87) and has potentially widespread utility in any situation where realistic rendering is possible.

Our approach is complementary to the long tradition of mechanistic modeling of sound localization. In contrast with mechanistic modeling, we do not produce specific hypotheses about underlying neural circuitry. However, the model gave rise to rich predictions of real-world behavior, and normative explanations of a large suite of perceptual phenomena. It should be possible to merge these two approaches, both by training model classes that are more faithful to biology (e.g. spiking neural networks) (88, 89), and by building in additional known biological structures to the neural network (e.g. replicating brainstem circuitry) (90).

### Limitations and Caveats

The strength of contemporary artificial neural networks lies in their ability to mediate some aspects of realistic behavior. We built on this strength to use them to reveal the behavioral characteristics of systems optimized under different conditions, which in some cases exhibit considerable behavioral similarities with human listeners. One limitation of this approach is that optimization of biological systems occurs in two distinct stages of evolution and development, which are not obviously mirrored in our machine optimization procedure. The procedure we used had separate stages of architectural selection and weight training, but these do not cleanly map onto evolution and development in biological systems. This limitation is shared by classical ideal observers, but limits the ability to predict effects that might be specific to one stage or the other, for instance involving plasticity (91).

Our model also shares many of the limitations that are common to supervised deep neural network models of the brain (92). The learning procedure is unlikely to have much in common with biological learning, both in the extent and nature of supervision (which involves millions of explicitly labeled examples) and in the learning algorithm, which is often argued to lack biological plausibility (88). The model class is also not fully consistent with biology, and so does not yield detailed predictions of neural circuitry.

The analogies with the brain thus seem most promising at the level of behavior and representations. Our results add to growing evidence that task-optimized models can produce human-like behavior for signals that are close to the manifold of natural sounds or images (43, 93). However, there is also evidence that artificial neural networks often exhibit substantial representational differences with humans, particularly for unnatural signals derived in various ways from a network (94–98), and it remains possible that the models here would exhibit similar divergences.

We chose to train models on a fixed representation of the ear. This choice was motivated by the assumption that the evolution of the ear was influenced by many different auditory tasks, such that it may not have been strongly influenced by the particular demands of sound localization, instead primarily serving as a constraint on biological solutions to the sound localization problem. However, the ear itself undoutedly reflects properties of the natural environment (99). It could thus be fruitful to “evolve” ears along with the rest of the auditory system, particularly in a framework with multiple tasks (43). Our cochlear model also does not replicate the fine details of cochlear physiology (100, 101) due to practical constraints of limited memory resources. These differences could in principle influence the results, although the similarity of the model results to those of humans suggests that the details of peripheral physiology beyond those that we modeled do not figure critically in the behavioral traits we examined.

The virtual world we used to train our models also no doubt differs in many ways from real-world acoustic environments. The rendering assumed point sources in space, which is inaccurate for many natural sound sources. The distribution of source locations was uniform relative to the listener, and both the listener and the sound sources were static, all of which are often not true of real-world conditions. And although the simulated reverberation replicated many aspects of real-world reverberation, it probably did not perfectly replicate the statistical properties of natural environmental impulse responses (102), or their distribution across environments. Our results suggest that the virtual world approximates the actual world in many of the respects that matter for spatial hearing, but the discrepancies with the real world could make a difference for some behaviors.

We also emphasize that despite presenting our approach as an alternative to ideal observer analysis (9), the resulting model almost surely differs in some respects from a fully ideal observer. The solutions reached by our approach are not provably optimal like classic ideal observers, and the model class and optimization methods could impose biases on the solutions. It is also likely that the architecture search was not extensive enough to find the best architectures for the task. Those caveats aside, the similarity to human behavior, along with the strong dependence on the training conditions, provides some confidence that the optimization procedure is succeeding to a degree that is scientifically useful.

### Future Directions

Our focus in this paper has been to model behavior, as there is a rich set of auditory localization behaviors for which normative explanations have traditionally been unavailable. However, it remains possible that the model we trained could be usefully compared to neural data. There is a large literature detailing binaural circuitry in the brainstem that could be compared to the internal responses of task-optimized models. The model could also be used to probe for functional organization in the auditory cortex, for instance by predicting brain responses using features from different model stages (38–43), potentially helping to reveal hierarchical stages of localization circuitry.

A model that can predict human behavior should also have useful applications. Our model showed some transfer of localization for specific sets of ears (Figure 5G), and could be used to make predictions about the extent to which sound rendering in virtual acoustic spaces (which may need to use a generic set of head-related transfer functions) should work for a particular listener. It can also predict which of a set of sound sources will be most compellingly localized, or worst localized (Figure 8). Such predictions could be valuable in enabling better virtual reality, or in synthesizing signals that humans cannot pinpoint in space.

The model we trained replicated human behavior in the most common traditional experimental setting, in which sound sources and the listener’s head are static. In real-world settings these assumptions are generally violated – sound sources often move relative to the listener, and the listener moves their head, often to better disambiguate front from back and more accurately localize (103). Our modeling approach could be straightforwardly expanded in this direction, with moving sound sources in the virtual training environment, and a network that can learn to move its head, potentially yielding explanations of auditory motion perception (104–106).

Another natural next step is to incorporate localization into auditory scene analysis, enabling localization of multiple concurrent sources as in the ‘cocktail party problem’ (107, 108). The model we built here has no natural way to report more than one sound source, but the virtual training approach could be naturally extended in this direction (109), potentially combining source separation or causal inference with localization, and helping to explain interactions between spatial hearing and source separation in humans (110–113). It will also be important to instantiate both recognition and localization in the same model, potentially yielding insight into the segregation of these functions in the brain (114). We envision extending the approach to encompass multi-modal perception (115), in which visual and auditory scene perception are jointly learned in a virtual world, enabling machine systems that more closely replicate human perceptual intelligence. More generally, the approach we take here – using deep learning to derive optimized solutions to perceptual or cognitive problems in different operating conditions – is broadly applicable to understanding the forces that shape complex, real-world, human behavior.

## Methods

### Training Data Generation

#### Virtual acoustic simulator - Image/Source method

We used a room simulator (45) to render Binaural Room Impulse Responses (BRIRs). This simulator used the image-source method, which approaches an exact solution to the wave equation if the walls are assumed to be rigid (50), as well as an extension to that method that allowed for more accurate calculation of the arrival time of a wave (116). This enabled the simulator to correctly render the relative timing between the signals received by the two simulated ears. Our specific implementation was identical to that used in (45), except for some custom optimization to take advantage of vectorized operations and parallel computation.

The room simulator operated in three separate stages. First, the image-source model calculated the positions of reflections of the source impulse forward in time for 0.5s. For each of these positions, the simulator placed an image symmetrically reflected about the wall of last contact. Second, the model accounted for the absorption spectra of the reflecting walls for each image location and filtered a broadband impulse sequentially using the absorption spectrum of the simulated wall material. Third, the room simulator found the direction of arrival for each image and convolved the filtered impulse with the head-related impulse response in the recorded set whose position was closest to the computed direction. This resulted in a left and right channel signal pair for each path from the source to the listener. Lastly, each of these signal pairs were summed together, factoring in both the delay from the time of arrival and the level attenuation based on total distance traveled by each reflection. The original authors of the simulator previously assessed this method’s validity and found that simulated BRIRs were good physical approximations to recorded BRIRs provided that sources were rendered more than one meter from the listener (45).

We used this simulator to render BRIRs for all elevations between 0° and 60° and all azimuths, spaced 5° in azimuth and 10° in elevation, at a distance of 1.4 meters, at each of a set of locations in 5 different rooms varying in size and material (listed below). This yielded 504 source positions per listener location. Listener locations were chosen subject to three constraints. First, the listener location had to be at least meters from the nearest wall (because sounds were rendered 1.4 meters from the listener). Second, the listener locations were located on a grid whose axes ran parallel to the walls of the room, with locations spaced 1 meter apart in each dimension. Third, the grid was centered on the center of the room. These constraints yielded 4 listener locations for the smallest room, and 81 listener locations for the largest room. This resulted in 71,064 pairs of BRIRs, each corresponding to a possible source-listener-room spatial configuration. Each BRIR took approximately 4 minutes to generate when parallelized across 16 cores. We parallelized generation of the full set of BRIRs across approximately 1000 cores on the MIT OpenMind Cluster, which allowed us to generate the full set of BRIRs in approximately 4 days.

| **Simulated Room Configurations** |  |
| --- | --- |
| Room Geometry (Length, Width, Height) | Room Material (Walls, Floor, Ceiling) |
| 9,9,10 | Plaster on Concrete, Carpet on Concrete, Acoustic Tiles 0.625" Thick |
| 5,4,2 | Brick, Carpet on Foam Padding, Plaster |
| 10,10,4 | Wood Paneling, Audience in Upholstered Seats, Sound Dampening Panels 1" Thick |
| 8,5,5 | Heavyweight Drapery, Carpet on Concrete, Plaster on Lath |
| 3,3,4 | Grass, No Reflections, No Reflections |

#### Virtual acoustic simulator - HRIRs used

The simulator relied on empirically derived Head Related Impulse Responses (HRIRs) to incorporate the effect of head shadowing and time delays without solving wave equations for the head and torso. Specifically, the simulator used a set of HRIRs recorded with KEMAR – a mannequin designed to replicate the acoustic effects of head and torso filtering on auditory signals. These recordings consisted of 710 positions ranging from −40° to +90° elevation at 1.4 meters (46).

#### Virtual acoustic simulator – Two-microphone array

For comparison with the networks trained with simulated ears, we also trained the same neural network architectures to localize sounds using audio recorded from a two microphone array. To train these networks, we simulated audio received from a two-microphone array by replacing each pair of HRIRs in the room simulator with a pair of fractional delay filters (i.e, that delayed the signal by a fraction of a sample). These filters consisted of 127 taps and were constructed via a sinc function windowed with a Blackman window, offset in time by the desired delay. Each pair of delay filters also incorporated signal attenuation from a distance according to the inverse square law, with the goal of replicating the acoustics of a two-microphone array. After substituting these filters for the HRIRs used in our main training procedure, we simulated a set of BRIRs as described above.

#### Natural sound sources

We collected a set of 455 natural sounds, each cut to two seconds in length. 300 of these sounds were drawn from a set used in previous work in the lab (117). Another 155 sounds were drawn from the BBC Sounds Effects Database, selected by the first author to be easily identifiable. The sounds included human and animal vocalizations, human actions (chopping, chewing, clapping etc.), machine sounds (cars, trains vaccums etc.) and nature sounds (thunder, insects, running water). All sounds were sampled at 44.1 kHz. Of this set, 385 sounds were used for training and another 70 sounds were withheld for model validation and testing. To augment the dataset, each of these was bandpass filtered with a two-octave-wide second-order Butterworth filter with center frequencies spaced in 1 octave steps starting from 100 Hz. This yielded 2,492 (2,110 training, 382 testing) sound sources in total. For each source, room, and listener location we randomly rendered each of the 504 positions with a probability 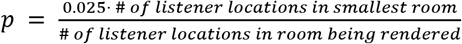. We used a base probability of 2.5% to limit the overall size of the training set and normalized by the number of listener locations in the room being used to render the current stimuus so that each room was represented equally in the dataset. This yielded 545,566 spatialized natural sound source stimuli in total (497,935 training, 47,631 testing).

#### Background noise sources

Background noise sources were synthesized using a previously described texture generation method that produced texture excerpts rated as highly realistic (118). The specific implementation of the synthesis algorithm was that used in (51). We used 50 different source textures obtained from in-lab collections (119). Textures were selected that synthesized successfully, both subjectively (sounding perceptually similar to the original texture) and objectively (the ratio between mean squared statistic values for the original texture and the mean squared error between the statistics of the synthesized and original texture was greater than 40dB SNR). We then rendered 1,000 5-second exemplars for each texture, cut to 2 seconds in length, for a total of 50,000 unique waveforms (1000 exemplars x 50 textures). Backgrounds were created by spatially rendering between 3 and 8 exemplars of the same texture at randomly chosen locations using the virtual acoustic simulator described above.

#### Generating training exemplars

For each training example the audio from one spatialized natural sound source and one spatialized background texture scene was combined (with a signal-to-noise-ratio sampled uniformly from 5 to 30 dB SNR) to create a single auditory scene that was used as a training example for the neural network. Each exemplar was passed through the cochlear model before being fed to the neural network.

#### Stimulus preprocessing for networks: Cochlear model

Training examples were pre-processed with a cochlear model to simulate the human auditory periphery. The output of the cochlear model is a time-frequency representation similar to a spectrogram, intended to represent the instantaneous mean firing rates in the auditory nerve. Cochleagrams were generated in a similar manner to previous work from our lab (118). However, the cochleagrams we used provided fine timing information to the network by passing rectified subbands of the signal instead of the envelopes of the subbands. This came at the cost of substantially increasing the dimensionality of the input relative to an envelope-based cochleagram.

The waveforms for the left and right channel were first upsampled to 48 kHz, then separately passed through a bank of 36 bandpass filters. These filters were regularly spaced on an equivalent rectangular bandwidth (ERB)_N_ scale (47) with bandwidths matched to those expected in a healthy human ear. Filter center frequencies ranged from 45 Hz to 16,975 Hz. Filters were zero-phase, with transfer functions in the frequency domain shaped as the positive portion of a cosine function. These filters perfectly tiled the frequency space such that the summed squared response of all filters was flat and allowed for reconstruction of the signal in that frequency range. Filtering was performed by multiplication in the frequency domain, yielding a set of subbands. The subbands were then transformed with a power function (0.3 exponent) to simulate the outer hair cells’ nonlinear compression. The results were then half-wave rectified to simulate the auditory nerve firing rates and were lowpass filtered with a cutoff frequency of 4 kHz to simulate the upper limit of phase locking in the auditory nerve (48), using a Kaiser-windowed sinc function with 4097 taps. All operations were performed in Python, but made heavy use of the NumPy and SciPy library optimization to decrease processing time. Code to generate cochleagrams in this way is available on the McDermott lab webpage (http://mcdermottlab.mit.edu).

In order to minimize artificial onset cues at the beginning and end of the cochleagram that would not be available to a human listener, we removed the first and last .35 seconds of the computed cochleagram and then randomly excerpted a 1-second segment from the remaining 1.3 seconds.

### Environment Modification for Unnatural Training Conditions

In each unnatural training condition, one aspect of the training environment was modified.

#### Anechoic environment

All echoes and reflections in this environment were removed. This was accomplished by setting the room material parameters for the walls, floor and ceiling to completely absorb all frequencies. This can be conceptualized as simulating a perfect anechoic chamber.

#### Noiseless environment

In this environment the background noise was removed by setting the SNR of the scene to 85dB. No other changes were made.

#### Unnatural sound sources

In this environment we replaced the natural sound sources with unnatural sounds consisting of repeating bandlimited noise bursts. For each 2 second sound source, we first generated a 200 ms 0.5 octave-wide noise burst with a 2 ms half-Hanning window at the onset and offset. We then repeated that noise burst separated by 200 ms of silence for the duration of the signal. The noise bursts in a given source signal always had the same center frequency. The center frequencies (the geometric mean of the upper and lower cutoffs) across the set of sounds were uniformly distributed on a log scale between 60 Hz and 16.8 kHz.

### Neural Network Models

The 36×16000×2 cochleagram representation was passed to a convolutional neural network (CNNs), which instantiated a feedforward, hierarchically organized set of linear and nonlinear operations. In our CNNs there were four different kinds of layers, each performing a distinct operation: (1) convolution with a set of filters, (2) a point-wise nonlinearity, (3) batch normalization, and (4) pooling. The first three types of layers always occurred in a fixed order (batch normalization, convolution, and a point-wise nonlinearity). We refer to a sequence of these three layers in this order as a “block”. Each block was followed by either another block or a pooling layer. Each network ended with either one or two fully connected layers feeding into the final classification layer. Below we define the operations of each type of layer.

#### Convolutional layer

A convolutional layer consists of a bank of K linear filters, each convolved with the input to produce K separate filter responses. Convolution performs the same operation at each point in the input, which in our case was the cochleagram. Convolution in time is natural for models of sensory systems as the input is a temporal sequence whose statistics are translation invariant. Convolution in frequency is less obviously natural, as translation invariance does not hold in frequency. However, approximate translation invariance holds locally in the frequency domain for many types of sound signals, and convolution in frequency is often present, implicitly or explicitly, in auditory models (120, 121). Moreover, imposing convolution greatly reduces the number of parameters to be learned, and we have found that neural network models train more readily when convolution in frequency is used, suggesting that it is a useful form of model regularization.

The input to a convolutional layer is a three-dimensional array with shape (n_in_,m_in_,d_in_) where n_in_ and m_in_ are the spectral and temporal dimensions of the input, respectively, and d_in_ is the number of filters. In the case of the first convolutional layer, n_in_ = 36 and m_in_ = 16,000, corresponding to the temporal and spectral dimensions of the cochleagram, and d_in_ = 2, corresponding to the left and right audio channels.

A convolution layer is defined by five parameters:

1. n_k_: The height of the convolutional kernels (i.e., their extent in the frequency dimension)
2. m_k_: The width of the convolutional kernels (i.e., their extent in the time dimension)
3. K: The number of different kernels
4. W: The kernel weights for each of the K kernels; this is an array of dimensions (n_k_, m_k_, d_in_, K).
5. B: The bias vector, of length K

For any input array X of shape (n_in_,m_in_,d_in_), the output of this convolutional layer is an array Y of shape (n_in_,m_in_ - m_k_ +1,K):

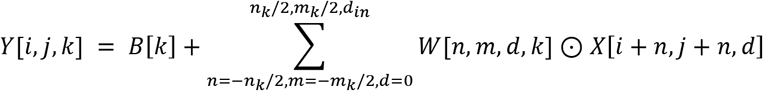

where i ranges from (1,…, n_in_), j ranges (1,…, m_in_), ⊙ represents pointwise array multiplication.

#### Boundary handling via valid padding in time

There are several common choices for boundary handling during convolution operations. In order to have the output of a convolution be the same dimensionality as the input, the input signal is typically padded with zeros. This approach - often termed ‘same’ convolution – has the downside of creating an artificial onset in the data that would not be present in continuous audio in the natural world, and that might influence the behavior of the network. To avoid this possibility, we used ‘valid’ convolution in the time dimension. This type of convolution only applies the filter at positions where every element of the kernel overlaps with the actual input. This eliminates artificial onsets at the start/end of the signal, but means that the output of the convolution will be slightly smaller than its input, as the filters cannot be centered over the first and last postions in the input without having part of the filter not overlap with the input data. We used ‘same’ convolution in the frequency dimension because the frequency dimension was much smaller than the time dimension, such that it seemed advantageous to preserve channels at each convolution stage.

#### Pointwise nonlinearity

If a neural network consists of only convolution layers, it can be mathematically reduced to a single matrix operation. A nonlinearity is needed for the neural network to learn more complex functions. We used rectified linear units (a common choice in current deep neural networks) that operate pointwise on every element in the input map according to a piecewise linear function:

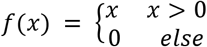

#### Normalization layer

The normalization layer applied batch normalization (122) in a pointwise manner to the input map. Specifically, for a batch B of training examples, consisting of examples {*X*_1_,…, *X*_*M*_}, with shape (n_in_,m_in_,d_in_), each example is normalized by the mean and variance of the batch:

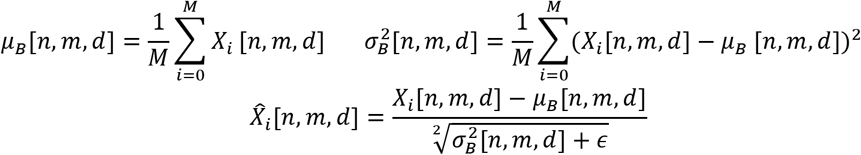

Where 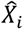 is the normalized three-dimensional matrix of the same shape as the input matrix and *ϵ* = 0.001 to prevent division by zero.

Throughout training the batch normalization layer maintains a cumulative mean and variance across all training examples, *μ*_*Total*_and 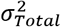. At test time 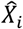 is calculated using *μ*_*Total*_ and 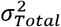 in place of *μ*_*B*_ and 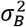.

#### Pooling layer

A pooling layer allows downstream layers to aggregate information across longer periods of time and wider bands of frequency. It downsamples its input by aggregating values across nearby time and frequency bins. We used max pooling, which is defined via 4 parameters:

1. p_h_, the height of the pooling kernel
2. p_w_, the width of the pooling kernel
3. s_h_, the stride in the vertical dimension
4. s_w_, the stride in the horizontal dimension

A pooling layer takes array X of shape (n_in_,m_in_,d_in_) and returns array Y with shape (n_in_/s_w_,m_in_/s_h_,d_in_) according to:

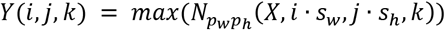

where 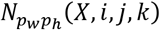 is a windowing function that takes a (p_w_, p_h_) excerpt of X of centered at (i,j) from filter k. The maximum is over all elements in the resulting excerpt.

#### Fully connected layer

A fully connected layer, also often called a dense layer, does not use the weight sharing found in convolutional layers, in which the same filter is applied to all positions within the input. Instead, each (input unit, output unit) pair has its own learned weight parameter and each output unit has its own bias parameter. Given input X with shape (n_in_,m_in_,d_in_), it produces output Y with shape (n_out_). It does so in two steps:

1. Flattens the input dimensions creating an input *X*_*flat*_ of shape (*n*_*in*_ ∙ *m*_*in*_ ∙ *d*_*in*_)
2. Multiplies *X*_*flat*_ by weight and bias matrices of shape (*n*_*out*_, *n*_*in*_ ∙ *m*_*in*_ ∙ *d*_*in*_) and (*n*_*out*_) respectively. This is implemented as:

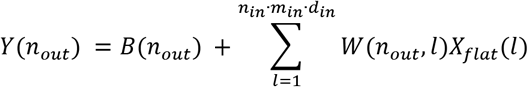

where *B*(*n*_*out*_) is the bias vector, *W*(*n*_*out*_, *l*) is the weight matrix, and *l* ranges from 1 to (*n*_*in*_ ∙ *m*_*in*_ ∙ *d*_*in*_) and indexes all positions in the flattened input matrix.

#### Softmax classifier

The final layer of every network was a classification layer, which consists of a fully connected layer where n_out_ is the number of class labels (in our case 504). The output of that fully connected layer was then passed through a normalized exponential function. Together this was implemented as:

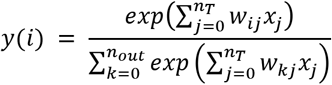

The vector y sums to 1 and all entries are greater than zero. This is often interpreted as a vector of label probabilities conditioned on the input.

#### Dropout during training

For each new batch of training data, dropout was applied to all fully connected layers of the network. Dropout consisted of randomly choosing 50% of the weights in the layer and temporarily setting them to zero, thus effectively not allowing the network access to the information at those positions. The other 50% of the weights were scaled up such that the expected value of the sum over all inputs was unchanged. This was implemented as:

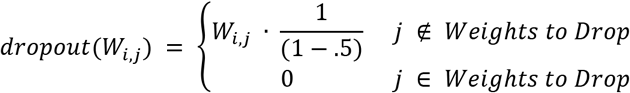

Dropout is common in neural network training and can be viewed as a form of model averaging where exponentially many models using different subsets of the input vector are being trained simultaneously (123). During evaluation dropout was turned off, so that all weights were used, and no weight scaling was performed.

### Neural Network Optimization

#### Architecture search - Overview

When neural networks are applied to a new problem it is common to use architectures that have previously produced good results on similar problems. However, most standard CNN architectures that operate on two-dimensional inputs have been designed for visual tasks and make assumptions based on the visual world. For example, most architectures assume that the units in the x and y dimension are equivalent, such that square filter kernels are a reasonable choice. However, in our problem the two input dimensions are not comparable (frequency vs time). Additionally, our input dimensionality is several orders of magnitude larger than standard visual stimuli (70k vs 1.1M), even though some relevant features occur on the scale of a few samples. For example, an ITD of 400 µs (a typical value) corresponds to only a 6 sample offset between channels. Given that our problem was distinct from many previous applications of standard neural network architectures, we performed an architecture search to find architectures that were well-suited to our task. First, we defined a space of architectures described by a small number of hyperparameters. Next, we defined discrete probability distributions for each hyperparameter. Lastly, we independently sampled from these hyperparameter distributions to generate architectures. We then trained each architecture for a brief period and selected the architectures that performed best on our task for further training.

#### Architecture search – Distribution over hyperparameters

To search over architectures we defined a space of possible architectures that were encoded via a set of hyperparameters. The space had the following constraints:

- There could be between 3 and 8 pooling layers for any given network.
- A pooling layer was preceded by between 1 and 3 blocks. Each block consisted of a batch norm, followed by a convolution, followed by a rectified linear unit.
- The number of channels (filters) in the network was always 32 in the first convolutional layer and in each successive convolutional layer could either double or remain the same.
- The penultimate stage of each network consisted of 1 or 2 fully connected layers containing 512 units each. Each of these was followed by a dropout layer.
- The final stage of each network was always a Softmax Classifier with 504 output units, corresponding to the 504 locations the network could report.

Discrete Prior Distributions for architecture search

- Number of pooling layers = [3,4,5,6,7,8]
- Number of blocks before each poolong layer = [1,2,3]

We picked the pooling and convolutional kernel parameters at each layer by uniformly sampling from the lists of values below. We chose these distributions to skew toward smaller values at deeper layers, approximately in line with the downsampling that resulted from pooling operations. Multiple copies of the same number increased the probability of that value being chose for the kernel size. Note that potential differences between the time and frequency dimensions of cochlear responses motivate the use of filters that are not square.

| **Network Layer** | **Convolutional Kernel width** | **Convolutional Kernel height** | **Pooling Kernel width** | **Pooling Kernel height** |
| --- | --- | --- | --- | --- |
| 1 | [4,8,16,32,64] | [1,2,3] | [2,4,8] | [1,2] |
| 2 | [4,8,16,32] | [1,2,3] | [2,4] | [1,2] |
| 3 | [2,4,8,16] | [1,2,3] | [2,4] | [1,2] |
| 4 | [2,4,8] | [1,2,3] | [1,2] | [1,2] |
| 5 | [2,4,8] | [1,2,3] | [1,2] | [1,1,2] |
| 6 | [2,3,4] | [1,2,3] | [1,1,2] | [1,1,2] |
| 7 | [2,3,4] | [1,2,3] | [1,1,1,2] | [1,1,1,2] |
| 8 | [2,3,4] | [1,2,3] | [1,1,1,2] | [1,1,1,2] |

#### Filter weight training

Throughout training, the parameters in each convolutional kernel and all weights from fully connected layers were iteratively adjusted to improve task accuracy via Mini-Batch Stochastic Gradient Descent (SGD) (124). Training was performed with 1.6 million sounds generated by looping over the 500,000 foreground sounds and combining each with a randomly selected background sound. Networks were assessed via a held out set of 50,000 test stimuli created by looping over the 48,000 validation set sound sources in the same manner. We trained it using a batch size of 16 and a Softmax Cross-Entropy loss function. The trainable weights in the convolutional layers and fully connected layers were updated using the gradient of the loss function, computed using backpropagation.

### Gradient checkpointing

The dimensionality of our input is sufficiently large (due to the high sampling rates needed to preserve the fine timing information in the simulated auditory periphery) as to preclude training neural networks using standard methodology. For example, consider training a standard network consisting of four max pooling layers (2 x 1 kernel), each preceded by one block. If there are 32 convolutional filters in the first layer, and double the number of filters in each successive layer, this network would require approximately 80GB of memory at peak usage, which exceeds the maximum memory of standard GPUs (currently varying between 12GB and 32GB). We addressed this problem using a previously proposed solution called gradient checkpointing (44).

In the standard backpropagation algorithm, we must retain the output from each layer of a network in memory because it is needed to calculate gradients for each updatable parameter. The gradient checkpointing algorithm we used trades speed for lower memory usage by not retaining each layer’s output during the forward pass, instead recomputing it a second time during the backward pass when gradients are computed. In the most extreme version, this would result in laboriously recomputing each layer starting with the original network input. Instead, the algorithm creates sparse, evenly spaced checkpoints throughout the network that save the output of selected layers. This strategy allows re-computation during backpropagation to start from one of these checkpoints, saving compute time. In practice, it also provides users with a parameter that allows them to select a speed/memory tradeoff that will maximize speed subject to a network fitting onto the available GPU. We created checkpoints at every pooling layer and found it kept our memory utilization below the 16GB limit of the hardware we used for all networks in the architecture search.

#### Network architecture selection and training

We performed our architecture search on the Department of Energy’s Summit Supercomputer at Oak Ridge National Lab. First, we randomly drew 1,500 architectures from our hyperparameter distribution. Next, we trained each architecture (i.e., optimized the weights of the convolutional and fully connected layers) using Mini-Batch Stochastic Gradient Descent for 15,000 steps, each with a batch size of 16, for a total of 240,000 unique examples, randomly drawn from the training set described above. We then evaluated the performance of each architecture on left-out data. This length of this training period was determined by the job limits on Summit; however, it was long enough to see significant reductions in the loss function for many networks. We considered the procedure adequate for architecture selection given that performance early in training is a good predictor of training performance late in training (125). In total, this architecture search took 2.05 GPU years and 45.2 CPU years.

We selected the ten best-performing architectures. They varied significantly, ranging from 4 to 6 pooling layers. We then retrained these 10 networks until a point where performance on the withheld validation set began to decrease, evaluating every 25,000 iterations. This occurred at 100,000 iterations for the normal, anechoic, and noiseless training conditions and at 150,000 iterations for the unnatural sounds training condition. Model architectures and the trained weights for each model are available online in the associated codebase: www.github.com/afrancl/BinauralLocalizationCNN.

### Real-World Evaluation

We tested the network in real-world conditions to verify generalization from the virtual training environment. We created a series of controlled spatial recordings in an actual conference room (part of our lab space, with dimensions distinct from the rooms in our virtual training environment) and then presented those to the trained network. We also made recordings of the same source sounds and environment with a two-microphone array to test the importance of naturally induced binaural cues (from the ears/head/torso).

#### Sound sources

We used 100 sound sources in total. 50 sounds sources were from our validation set of withheld environmental sounds, and the remaining 50 sound sources were taken from the GRID dataset of spoken sentences (126). For the examples from the GRID dataset, we used 5 sentences from each of 10 speakers (5 male and 5 female). The network performed similarly for stimuli from the GRID dataset as for our validation set stimuli. All source signals were normalized to the same peak amplitude before the recordings were made.

#### Recording setup

We made the first set of recordings using a KEMAR head and torso simulator mannequin built by Knowles Electronics to replicate the shape and absorbency of a human head, upper body, and pinna. The KEMAR mannequin contains a microphone in each ear, recording audio similar to that which a human would hear in natural conditions. The audio from these microphones was then passed through Etymotic Research preamplifiers designed for the KEMAR mannequin before being passed to the Zoom 8 USB to Audio Converter. Finally it was passed to Audacity where the left and right channel were simultaneously recorded at 48kHz.

We made recordings of all 100 sounds at every azimuth (relative to the KEMAR mannequin) from 0° to 360° in 30° increments. This led to a total set of 1,200 recordings in total. All source sounds were played meters from the vertical axis of the mannequin using a KRK ROKIT 7 speaker. The audio was played using Audacity and converted to analogue signal using a Zoom 8 USB to Audio Converter.

Recordings were made in our main lab space in building 46 on the MIT campus, which is roughly 7×6×3 meters. The room is filled with furniture, shelves and has multiple windows and doors (Figure 2A). This setup was substantially different from any of the simulated rooms in the virtual training environment, in which all rooms were convex, empty, and had smooth walls. During the recordings there was low-level background noise from the HVAC system, the refrigerator, and lab members talking in surrounding offices. For all recordings the mannequin was seated in an office chair, with the head approximately 1 meter from the ground.

#### Two-microphone array baseline

We made the second set of recordings using the same sound sources, room, and recording equipment as above, but with the KEMAR mannequin replaced with a 2-microphone array consisting of two Beyerdynamic MM-1 Omnidirectional Microphones separated by 15cm (the same distance separating the two microphones in the mannequin ears). The microphone array was also elevated approximately 1 meter from the floor using a microphone stand.

#### Baseline algorithms

We evaluated our trained neural networks against a variety of baseline algorithms. These include: Steered-Response Power Phase Transform (SRP) (57), Multiple Signal Classification (MUSIC) (56), Coherent Signal-Subspace Method (CSSM) (55), Weighted Average of Signal Subspaces (WAVES) (58), and Test of Orthogonality of Projected Subspaces (TOPS) (59). In each case we used the previously validated and published algorithm implementations in Pyroomacoustics (127). We also created a baseline neural network trained using a simulation of the two-microphone array described in the previous section within the virtual training environment.

The results shown in Figure 2F for the baselines (aside from our two-microphone array baseline network) all plot localization of the KEMAR mannequin recordings. We found empirically that the baseline methods performed better for the KEMAR recordings than for the two-microphone array recordings, presumably because the mannequin head increases the effective distance between the microphones. The baseline algorithms require prior knowledge of the inter-microphone distance. In order to make the baselines as strong as possible relative to our method, we searched over all distances less than 50cm and found that an assumed distance of 26cm yielded the best performance. We then evaluated the baselines at that assumed distance. This optimal assumed distance is greater than the actual inter-microphone distance of 15cm, consistent with the idea that the mannequin head increases the effective distance between microphones.

### Simulated Experiments

#### Overview

We simulated a suite of classic psychoacoustic experiments on the 10 trained neural networks, using the same stimuli for each network. We then calculated the mean response across networks for each experimental condition and calculated error bars by bootstrapping across the 10 networks. This approach can be interpreted as marginalizing out uncertainty over architectures in a situation in which there is no single obviously optimal architecture (and where the space of architectures is so large that it is probably not possible to find the optimum even if it exists). Moreover, recent work suggests that internal representations across different networks trained on the same task can vary considerably (49), so this approach aided in mitigating the individual idiosyncrasies of any given network. The approach could also be viewed as treating every network as an individual experimental participant, calculating means and error bars as one would in a standard human psychophysics experiment.

In each experiment, stimuli were run through our cochlear model and passed to the network, whose localization responses were recorded for each stimulus.

#### Sensitivity to interaural time and level differences – Stimuli

We reproduced the experimental stimuli from (60), in which ITDs and ILDs were added to 3D spatially rendered sounds. In the original experiment, participants stood in a dark anechoic room and were played spatially rendered stimuli with modified ITDs or ILDs via a set of headphones. After each stimulus presentation, participants oriented their head towards the percieved location of the stimulus and pressed a button. The experiment included 13 participants (5 male) ranging in age from 18-35 years old.

Stimulus generation for the model experiment was identical to that in the original experiments apart from using our acoustic simulator to render the sounds. First, we generated highpass and lowpass noise bursts with passbands of 4-16 kHz and 0.5-2 kHz, respectively (44.1 kHz sampling rate). Each noise burst was 100 ms long with a 1 ms squared-cosine ramp at the beginning and end of the stimulus. We randomly jittered the starting time of the noise burst by padding the signal to 2,000 ms in total length, constrained such that the entire noise burst was contained in the middle second of the 2s audio signal. These signals were then rendered at 0° elevation, with azimuth varied from 0°-355° (in 5° steps) for a total of 72 locations. All signals were rendered using our virtual acoustic simulator in an anechoic environment without any background noise.

Next, we created versions of each signal with an added ITD or ILD bias. ITD biases were ±300 µs and ±600 µs and ILD biases were ±10 dB and ±20 dB (Figure 3A). As in the original publication (60), we prevented presentation of stimuli outside the physiological range by restricting the 400µs/10dB biases to signals rendered less than 40° away from the midline and restricting the 600µs/20dB biases to signals rendered less than 20° away from the midline. In total there were 4 stimulus sets (2 passbands x 2 types of bias) of 266 stimuli (72 locations with no bias, 52 locations at ± medium bias, 45 locations ± large bias). We replicated the above process 20 times with different exemplars of bandpassed noise, increasing each stimulus set size to 5,320 (20 exemplars of 266 stimuli).

#### Sensitivity to interaural time and level differences – Data analysis

We measured the perceptual bias induced by the added ITD or ILD bias in the same manner as the published analysis of human listeners (60).

We first calculated the naturally-occurring ITD and ILD for each sound source position (varying in azimuth, at 0° elevation) from the HRTFs used to train our networks. For ITDs, we ran the HRTFs for a source position through our cochlear model and found the ITD by cross-correlating the cochlear channels whose center frequency was closest to 600, 700 and 800 Hz and taking the median ITD from the three channels. For ILDs, we computed power spectral density estimates via Welch’s method (29 samples per window, 50% overlap, Hamming windowed) for each of the two HRTFs for a source position and integrated across frequencies in the stimulus passband. We expressed the ILD as the ratio between the energy in the left and right channel in decibels, with positive values corresponding to more power in the right ear. This set of natural ILDs and ITDs allowed us to map the judged position onto a corresponding ITD/ILD.

For each stimulus with added ITD, we used the response mapping described above to calculate the ITD of the judged source position. Next, we calculated the ITD for the judged position of the unaltered stimulus using the same response mapping. The perceptual effect of the added ITD was calculated as the difference between these two ITD values, quantifying (in microseconds) how much the added stimulus bias changed the response of the model. The results graphs plot the added stimulus bias on the x-axis and response bias on the y-axis. The slope of the best-fitting regression line (the ‘Bias Weight’ shown in the subplots of Figure 3B&C) provides a unitless measure of the extent to which the added bias affects the judged position. We repeated an analogous process for ILD bias using the natural ILD response mapping, yielding the bias in decibels. The graphs in Figure 3D plot the mean response across the 10 networks with standard error of the mean (SEM) computed via bootstrap over networks.

#### Azimuthal localization of broadband sounds – Stimuli

We reproduced the stimulus generation from (68). In their experiment, participants were played 6 broadband white noise bursts, with 3 bursts played from a reference speaker followed by 3 noise bursts played from one of two target speakers, located 15° to the left or right of the reference speaker. Participants reported whether the latter 3 noise bursts were played from the left or right target speaker. The experiment included 16 participants between the ages of 18 and 35 years old.

We measured network localization performance using the same stimuli as in the original paper, but rendered the stimulus at a single location and measured performance with an absolute, instead of relative, localization task. The stimuli presented to the networks consisted of 3 pulses of broadband white noise. Each noise pulse was 15 ms in duration and the delay between pulses was 100 ms. A 5 ms cosine ramp was applied to the beginning and end of each pulse. We generated 100 exemplars of this stimulus using different samples of white noise (44.1 kHz sampling rate). The stimuli were zero-padded to 2,000 ms in length, with the temporal offset of the three-burst sequence randomly sampled from a uniform distribution such that all three noise bursts were fully contained in the middle second of audio. We then rendered all 100 stimuli at 0° elevation and azimuthal positions ranging from 0° to 355° in 5° steps. All stimuli were rendered in an anechoic environment without any background noise using our virtual acoustic simulator. This led to 5,200 stimuli in total (100 exemplars at each of 52 locations).

#### Azimuthal localization of broadband sounds – Data analysis

Because human participants in the analogous experiment judged relative position in the frontal hemifield, prior to calculating the model’s accuracy we eliminated front-back confusions by mirroring model responses of each stimulus across the coronal plane. For example, the 10° and 170° positions were considered equivalent. We then calculated the difference in degrees between the rendered azimuthal position and the position judged by the model. We calculated the mean error for each rendered azimuth for each network. The graph in Figure 4C plots the mean error across networks. Error bars are SEM, bootstrapped over networks.

#### Integration across frequency – Stimuli

We reproduced stimuli from (69). In the original experiment, human participants were played a single noise burst, varying in bandwidth and center frequency, from one of 8 speakers spaced 15° in azimuth. Participants judged which speaker the noise burst was played from. The experimenters then calculated the localization error in degrees for each bandwidth and center frequency condition. The experiment included 33 participants (26 female) between the ages of 18 and 36 years old.

The stimuli varied in bandwidth (pure tones, and noise bursts with bandwidths of 1/20, 1/10, 1/6, 1/3, 1, and 2 octaves wide; all with 44.1 kHz sampling rate). All sounds were 200 ms long with a 20 ms squared-cosine ramp at the beginning and end of the sound. All pure tones had random phase. All other sounds were bandpass-filtered white noise with the geometric mean of the passband cutoffs at 250, 2,000, or 4,000 Hz (as in (69)).

For the model experiment, the stimuli were zero-padded to 2000ms in length, with the temporal offset of the noise burst randomly sampled from a uniform distribution such that the noise burst was fully contained in the middle second of audio. We generated 30 exemplars of each bandwidth/frequency pair using different exemplars of white noise or random phase for noise and tones, respectively. Next, we rendered all stimuli at 0° elevation and azimuthal positions ranging from 0° to 355° in 5° steps. All stimuli were rendered in an anechoic environment without any background noise using our virtual acoustic simulator. This led to 45,360 stimuli in total (30 exemplars x 72 positions x 3 center frequencies x 7 bandwidths).

#### Integration across frequency – Data analysis

Because human participants in the original experiment judged position in the frontal hemifield, prior to calculating the model’s accuracy we again eliminated front-back confusions by mirroring model responses of each stimulus across the coronal plane. We then calculated the difference in degrees between the rendered azimuthal position and the azimuthal position judged by the model. For each network we calculated the root-mean-squared error for each bandwidth. The graph in Figure 4F plots the mean of this quantity across networks. Error bars are SEM, bootstrapped over networks.

#### Use of ear-specific cues to elevation – Stimuli

We simulated a change of ears for our networks, analogous to the ear mold manipulation in (72)). In the original experiment (72), participants sat in a dark anechoic room and were played broadband white noise bursts from a speaker on a robotic arm that moved ±30° in azimuth and elevation. Participants reported the location of each noise burst by saccading to the perceived location. After collecting a baseline set of measurements, participants were fitted with plastic ear molds (Figure 5A), which modified the location-dependent filtering of their pinnae, and performed the same localization task a second time. The experimenters plotted the mean judged location for each actual location before and after fitting subjects with plastic ear models (Figure 5B&C). The experiment included 4 participants between the ages of 22 and 44 years old.

For the model experiment, instead of ear molds we substituted HRTFs from the CIPIC dataset (128). The CIPIC dataset contains 45 sets of HRTFs, each of which is sampled at azimuths from −80 to +80 in 25 steps of varying size, and elevations from 0 to 360 in 50 steps of varying size. We generated 500 ms broadband (0.2-20 kHz) noise bursts sampled at 44.1 kHz (as in (72)). We then zero-padded these sounds to 2,000 ms, with the temporal offset of the noise burst randomly sampled from a uniform distribution such that it was fully contained in the middle second of audio. We generated 20 such exemplars using different samples of white noise. We then rendered each stimulus at ±20 and ±10° azimuths and 0°, 10°, 20°, and 30° elevation for all 45 sets of HRTFs as well as the standard set of HRTFs (i.e., the one used for training the networks). This led to a total of 14,720 stimuli (46 HRTFs x 4 azimuths x 4 elevations). The rendered locations were slightly different from those used in (72) as we were constrained by the locations that were measured for the CIPIC dataset.

#### Use of ear-specific cues to elevation – Data analysis

The results graphs for this experiment (Figure 5B-E) plot the judged source position for each of a set of rendered source positions, either for humans (Figure 5B&C) or the model (Figure 5D&E). For the model results, we first calculated the mean judged position for each network for all stimuli rendered at each source position. The graphs plot the mean of this quantity across networks. Error bars are the SEM, bootstrapped over networks. In Figure 5D we plot model responses for stimuli rendered using the HRTFs used during network training. In Figure 5E we plot the average model responses for stimuli rendered with 45 sets of HRTFs from the CIPIC database (none of which were used during network training). In Figures 5F&G we plot the results separately for each alternative set of HRTFs, averaged across elevation or azimuth. The thickest bolded line denotes the mean performance across all HRTFs, and thinner bolded lines denote HRTFs at the 5^th^, 25^th^, 75^th^, and 95^th^ percentiles order by error. Each line plots the mean over the 10 networks.

#### Limited spectral resolution of elevation cues – Stimuli

We ran a modified version of the spectral smoothing experiment in (74) on our networks using the training HRTFs. The original experiment (74) measured the effect of spectral detail on human sound localization. The experimenters first measured HRTFs for 4 participants. Participants then sat in an anechoic chamber and were played broadband white noise bursts presented in one of two ways. The noise burst was either played directly from a speaker in the room or virtually rendered at the position of the speaker using the participant’s HRTF and played from a set of open backed earphones worn by the participant. The experimenters manipulated the spectral detail of the HRTFs as described below. On each trial, two noise bursts (one each played via the two presentation methods) were played in random order and participants judged which of the two noise bursts were played via earphones. In practice this judgment was performed by noticing changes in the apparent sound position that occurred when the HRTFs were sufficiently degraded. The results of the experiment were expressed as the accuracy in discriminating between the two modes of presentation as a function of the amount of spectral detail removed (Figure 5I). The experiment included 4 participants.

The HRTF is obtained from the Fourier transform of the head-related impulse response (HRIR), and thus can be expressed as:

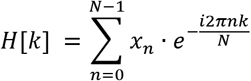

where x is the head-related impulse response, N is the number of samples in the HRTF and k = [0,N-1]. To smooth the HRTF, we first compute the log-magnitude of H[k].. This log-magnitude of the HRTF can be decomposed into frequency components via the discrete cosine transform:

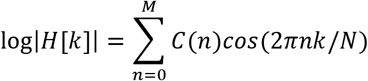

where C(n) is the nth cosine coefficient of log |*H*[*k*]| and M = N/2.

As in (74), we smoothed the HRTF by reconstructing it with M < N/2. In the most extreme case where M=0, the magnitude spectrum was perfectly flat at the average value of the HRTF. Increasing M increases the number of cosines used for reconstruction, leading to more spectral detail (Figure 5H). After smoothing, we calculated the minimum phase filter from the smoothed magnitude spectrum, adding a frequency-independent time delay consistent with the original HRIR. Our HRIRs consisted of 512 time points, corresponding to a maximum of 256 points in its cosine series.

We repeated this smoothing process for each left and right HRTF at each spatial position. We then generated 20 exemplars of broadband white noise (0.2-20kHz, 2000 ms length) with a 10 ms cosine ramp at the beginning and end of the signal. Each exemplar was rendered at 0° elevation and azimuthal positions ranging from 0° to 355° in 5° steps using each smoothed set of HRTFs. This yielded 12,960 stimuli (9 smoothed sets of HRTFS x 20 exemplars x 72 locations).

#### Limited spectral resolution of elevation cues – Data analysis

For the model, the effect of the smoothing was measured as the difference in degrees between the judged position and the rendered position for each stimulus. Figure 5J plots the mean error across networks for each smoothed set of HRTFs. Error bars are SEM, bootstrapped over networks. Figures 5K&L plots the mean judged azimuth (left) and elevation (right) vs. the actual rendered azimuth and elevation, plotted separately for each smoothing level. Each line is the mean response pooled across networks. Error bars are shown as bands around the line and show SEM, bootstrapped over networks.

#### Precedence effect – Stimuli

For the basic demo of the precedence effect (Figure 6B) we generated a click consisting of a single sample at +1 surrounded by zeros. We then rendered that click at ±45 azimuth and 0° elevation in an anechoic room without background noise using our virtual acoustic simulator. We added these two rendered signals together, temporally offsetting the −45° click behind the 45° click by an amount ranging from 1 to 50 ms. We then zero-padded the signal to 2000 ms, sampled at 44.1 kHz, and randomly varied the temporal offset of the click sequence, constrained such that all nonzero samples occurred in the middle second of the stimulus. For each delay value, we created 100 exemplars with different start times.

To quantitatively compare the precedence effect in our model with that in human participants, we reproduced the stimuli from (80). In the original experiment, participants were played two broadband pink noise bursts from two different locations. The leading noise burst came from one of 6 locations (±20°, ±40°, or ±60°) and the lagging noise burst came from 0°. The lagging noise burst was delayed relative to the leading noise burst by 5, 10, 25, 50, or 100 ms. For each pair of noise bursts, participants reported whether they perceived one or two sounds and the judged location for each perceived sound. The experimenters then calculated the mean localization error separately for the leading and lagging click for each time delay (Figure 6C). The experiment included 10 participants (all female) between the ages of 19 and 26 years old.

For both the human and network experiments, stimuli were 25 ms pink noise bursts, sampled at 44.1kHz, with a 2 ms cosine ramp at the beginning and end of the burst. For the network experiment, we generated two stimuli for each pair of noise burst positions, one where the 0° noise burst was the lead click and another where it was the lag click. For each delay value, location and burst order, we created 100 exemplars with different start times. Stimuli were delayed and padded as described above.

#### Precedence effect – Data analysis

Because human experiments on the precedence effect typically query participants about positions in the frontal hemifield, we corrected for front-back confusions by mirroring model responses of each stimulus across the coronal plane. Figure 6B plots the mean judged position at each inter-click delay, averaged across the means of the 10 individual networks. Error bars are SEM, bootstrapped over networks.

To generate Figure 6D (plotting the results of the model version of the Litovsky and Godar experiment) we calculated errors for each stimulus between the model’s judged position and the positions of the leading and lagging clicks. We calculated the average lead click error and average lag click error for each network at each delay. Figure 6D plots the mean across the mean error for each network. Error bars are SEM, bootstrapped over networks.

#### Instrument note localization – Stimuli

To assess the ability of the model to predict localization behavior for natural sounds, we rendered a set of instruments playing notes at different spatial positions. Instruments were sourced from the Nsynth Dataset (81), which contains a large number of musical notes from a wide variety of instruments. We used the validation set, which contained 12,678 notes sampled from 53 instruments. For each note, room in our virtual environment, and listener location within each room, we randomly rendered each of the 72 positions (0° elevation, 0°-355° azimuth in 5° steps) with a probability 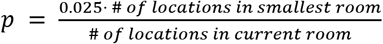.

We used a base probability of 2.5% to limit the overall size of the test set and normalized by the number of locations in the current room so that each room is represented equally in the test set. This yielded a total of 456,580 stimuli.

#### Instrument note localization – Data analysis

We anticipated performing a human instrument note localization experiment in an environment with speakers in the frontal hemifield, so we corrected for front-back confusions by mirroring model responses of each stimulus across the coronal plane. Different instruments in the dataset contained different subsets of pitches. To ensure that differences in localization accuracy would not be driven solely by the instrument’s pitch range, we limited analysis to instruments for which the dataset contained all notes in the octave around middle C (MIDI note 55 through 66) and performed all analysis on notes in that range. This yielded 43 instruments and 1860 unique notes. We calculated the mean localization error for each network judgment by calculating the absolute difference, in degrees, between the judged and rendered location. We then averaged the error across networks and calculated the mean error for each of the 1860 remaining notes from the original dataset. We plotted the distributions of the mean error over notes for each instrument (8A).

To characterize the density of the spectrum we computed its spectral flatness. We first estimated the power spectrum *x*(*n*) using Welch’s method (window size of 2000 samples, 50% overlap). The spectral flatness was computed for each note of each instrument as:

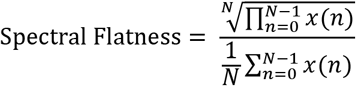

We averaged the spectral flatness across all notes of an instrument and then computed the Spearman correlation of this measure with the network’s mean accuracy for that instrument.

### Analysis of Results of Alternative Training Conditions

#### Human-model dissimilarity

We assessed the effect of training condition on model behavior by quantifying the extent of the dissimilarity between the model psychophysical results and the human results. For each results graph, we measured human-model dissimilarity as the root-mean-squared error between corresponding y-axis values in the human and model experiments. In order to compare results between experiments, before measuring this error, we min-max normalized the y-axis to range from 0 to 1. For experiments with the same y-axis for human and model results, we normalized the model and human data together (i.e., taking the min and max values from the pooled results). For experiments where the y-axes were different for human and model results (because the tasks were different, as in Figures 4B&C and 5I&J), we normalized the data individually for human and model results.

The one exception was the Ear Alteration experiment (Figure 5A-G), in which the result of primary interest was the change in judged location relative to the rendered location, and for which the locations were different in the human and model experiments (due to constraints of the HRTF sets that we used). To measure the human-model dissimilarity for this experiment, we calculated the error between the judged and rendered location for each point on the graph, for humans and the model. We then calculated human-model dissimilarity between these error values, treating the two grids of locations as equivalent. This approach would fail to capture some patterns of errors but was sufficient to capture the main effects of preserved azimuthal localization along with the collapse of elevation localization.

This procedure yielded dissimilarity measure that varied between 0 and 1 for each experiment, where 0 represents a perfect fit to the human results. For Figure 7B, we then calculated the mean of this dissimilarity measure over the seven experiments. To generate error bars, we bootstrapped across the 10 networks and recalculated all results graphs and the corresponding mean normalized error for each bootstrap sample. Error bars in Figure 7B plot the SEM of this distribution. Additionally, we plotted the mean normalized error individually for each of the 10 networks (Supplemental Figure 4).

#### Cohen’s D

To assess how training conditions impacted individual psychophysical effects, we measured the effect size of the difference between human-model divergence in the normal and alternative training conditions for each psychophysical effect. Specifically, we measured Cohen’s D for each experiment:

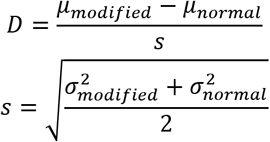

where *μ* and *σ*^2^ are the mean and variance of the human-model divergence across our 10 networks for the normal or modified training condition. We calculated error bars on Cohen’s D by bootstrapping across the 10 networks, computing the effect size for each bootstrap sample. Figure 7C plots the mean and SEM of this distribution.

### Statistics

#### Psychophysical experiments

For plots comparing models on real world data (Figure 2F and 2G), error bars are SEM, bootstrapped over stimuli and networks (because there was only one version of the baseline methods). For plots assessing duplex theory (Fig 3C), azimuth sensitivity (Figure 4C), bandwidth sensitivity (Figure 4F), changing ears (Figure 5D and 5E), spectral sensitivity (Figure 5J), and precedence effect (Figure 6B and 6D) error bars are SEM, bootstrapped across networks.

To assess the significance of the interaction between the stimulus frequency range and the magnitude of the ITD/ILD bias weights (Figure 3C), we calculated the difference of differences in bias weights across the 4 stimulus/cue-type conditions:

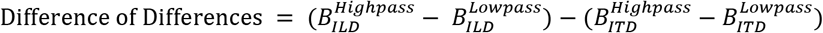

where B denotes the bias weight for each condition). We calculated the difference of differences bootstrapped across models with 10,000 samples. As this difference of differences exceeded 0 for all 10,000 bootstrap samples, we fit a Gaussian distribution to the histogram of values for the 10,000 bootstrap samples and calculated the p-value for a value of 0 or smaller from the fitted Gaussian.

We assessed the significance of the lowpass ILD bias weight (Figure 3C) by bootstrapping across networks, again fitting a Gaussian distribution to the histogram of bias weights from each bootstrap sample and calculating the p-value for a value of 0 or smaller from the fitted Gaussian.

#### Statistical significance of individual training conditions

We assessed the statistical significance of the effect of individual training conditions by bootstrapping the normalized error measure described above across networks for the natural training condition (Figure 7B).

We fit a Gaussian distribution to the histogram of the normalized error for each bootstrap sample and calculated the p-value for the mean normalized error measure for each modified training condition under the fitted Gaussian.

We also assessed the statistical significance of the effect size of the change to individual experiment results relative to other experiments when training in alternative conditions (Figure 7C). First we measured Cohen’s D across networks as described above for 10,000 bootstrap samples of the 10 networks, leading to a separate distribution for each experiment. For each experiment of interest, we assessed the probability that a value at or below the mean Cohen’s D of each other experiment could have occurred under the bootstrap distribution. The histogram of bootstrap samples was non-Gaussian so we calculated this probability by counting the number of values at or below the mean for each condition and reported the proportion of such values as the p-value.

We assessed the statistical significance of the effect of training condition on real-world localization performance (Figure 7E) by bootstrapping the RMS localization error across networks. We fit a Gaussian distribution to the histogram of RMS error for the normal training condition and calculated the probability that a value could have been drawn from that Gaussian at or above the mean RMS error for each alternative training condition.

## Acknowledgements

The authors thank Oak Ridge National Laboratories for use of their SUMMIT supercomputing facility, John Cohn for assistance securing computing resources, Bill Yost for sharing data, Ray Gonzalez for writing the cochleagram generation codebase, Jenelle Feather and Mark Saddler for their support and technical advice, members of the McDermott lab for their comments on the manuscript (especially Dana Boebinger, Maddie Cusimano, Jenelle Feather, Malinda McPherson, Mark Saddler, and James Traer), and Steve Colburn for lending us his KEMAR mannequin and for advice throughout the project.

## Notes

### Competing Interest Statement

The authors have declared no competing interest.

