## Supplemental Figures and Tables for "Deep neural network models of sound localization reveal how perception is adapted to real-world environments"

### Supplemental Information

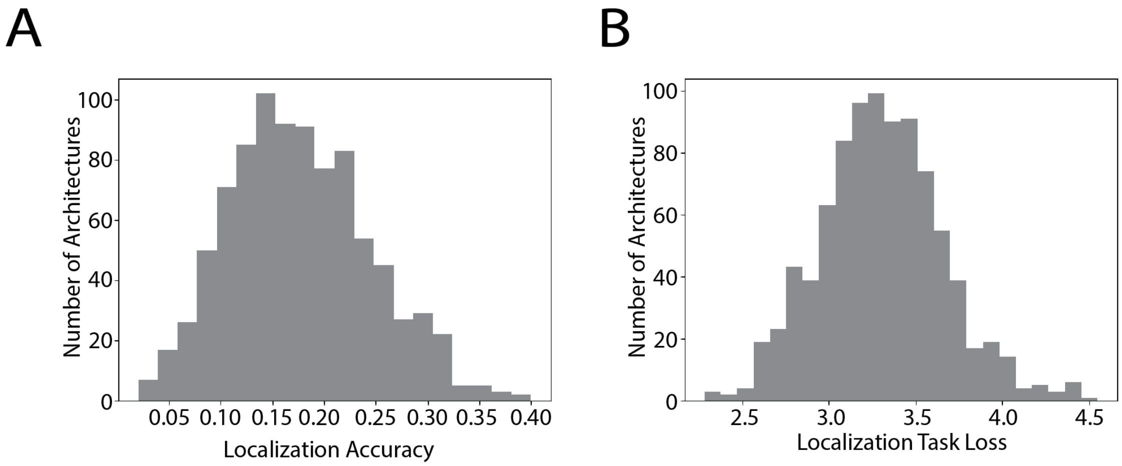


**Supplemental Figure 1.** A. Histogram of validation set accuracies for neural network architectures after 15k steps of training during architecture search. B. Histogram of validation set losses for neural network architectures after 15k steps of training during architecture search.

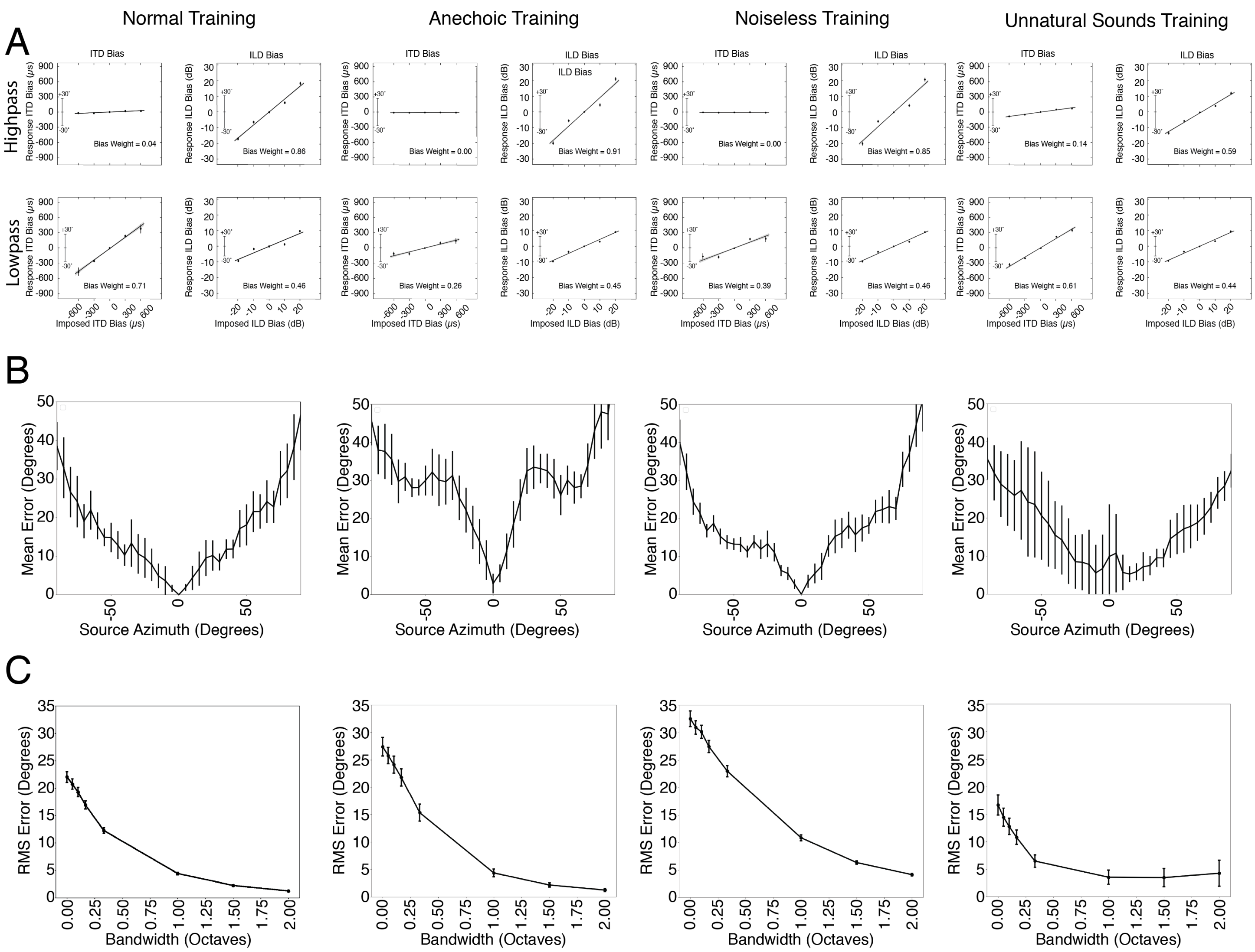


**Supplemental Figure 2.** Model psychophysics across training conditions for first three psychophysical experiments. A. Model sensitivity to interaural time and level differences (Figure 3D). B. Model accuracy for broadband noise at different azimuthal positions (Figure 4C). C. Effect of bandwidth on model localization of noise bursts (Figure 4F)

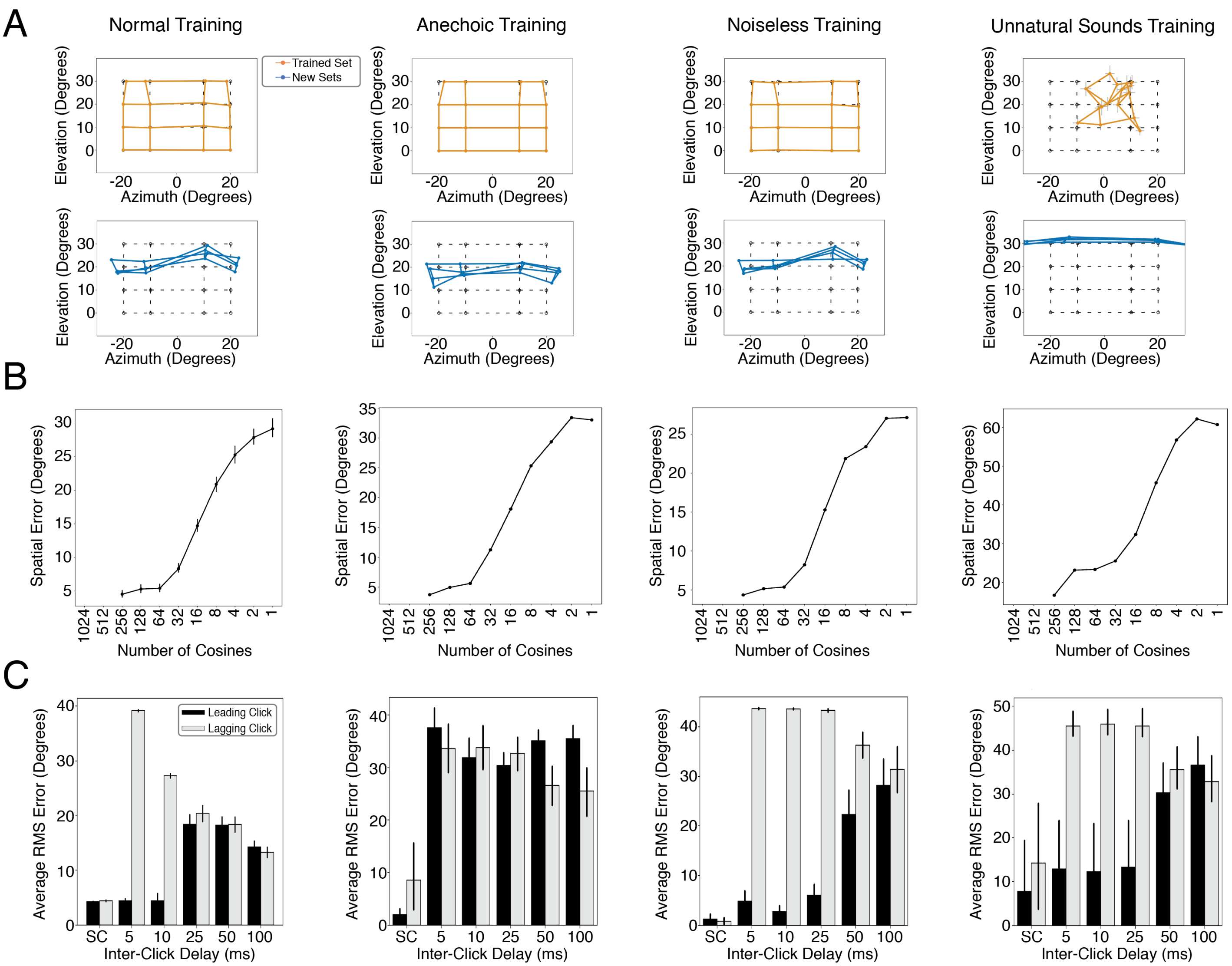


**Supplemental Figure 3.** Model psychophysics across training conditions for last three psychophysical experiments. A. Sound localization by the model in azimuth and elevation before and after ear alteration (Figure 5 D&E). B. Effect of spectral smoothing on model sound localization accuracy (Figure 5J). C. Model error in localization of the leading and lagging clicks in the precedence effect experiment, as a function of delay (Figure 6D).

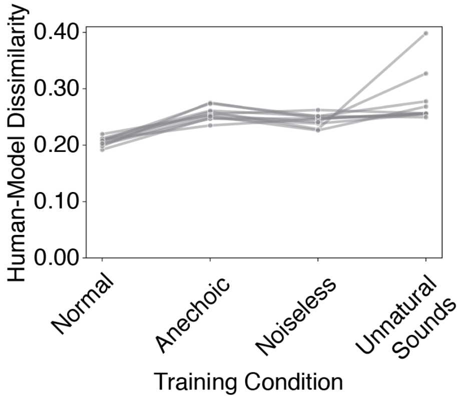


**Supplemental Figure 4.** Human-model dissimilarity for natural and unnatural training conditions for individual neural networks.

**Supplemental Table 1.** Summary of network architectures

| | Network Architecture Numbers | | | | | | | | | |
|---|---|---|---|---|---|---|---|---|---|---|
| Operation | 1 | 2 | 3 | 4 | 5 | 6 | 7 | 8 | 9 | 10 |
| 1 | Conv[1,8,32] | Conv[2,8,32] | Conv[1,4,32] | Conv[3,8,32] | Conv[2,32,32] | Conv[1,64,32] | Conv[1,16,32] | Conv[1,64,32] | Conv[3,32,32] | Conv[2,4,32] |
| 2 | Relu | Relu | Relu | Relu | Pool[1,2] | Pool[1,8] | Relu | Relu | Relu | Pool[2,2] |
| 3 | Bn | Bn | Bn | Bn | Relu | Relu | Bn | Bn | Bn | Relu |
| 4 | Conv[1,64,32] | Conv[3,16,32] | Conv[1,32,32] | Conv[3,8,32] | Bn | Bn | Conv[1,8,32] | Conv[2,16,32] | Conv[2,16,32] | Bn |
| 5 | Relu | Relu | Pool[1,8] | Pool[1,2] | Conv[1,4,64] | Conv[2,4,64] | Pool[1,2] | Pool[1,8] | Pool[1,4] | Conv[2,4,32] |
| 6 | Bn | Bn | Relu | Relu | Pool[1,4] | Relu | Relu | Relu | Relu | Pool[1,4] |
| 7 | Conv[1,64,32] | Conv[2,4,32] | Bn | Bn | Relu | Bn | Bn | Bn | Bn | Relu |
| 8 | Pool[1,8] | Pool[1,8] | Conv[3,32,64] | Conv[1,32,64] | Bn | Conv[1,32,64] | Conv[2,4,64] | Conv[2,4,64] | Conv[2,32,64] | Bn |
| 9 | Relu | Relu | Relu | Relu | Conv[3,2,64] | Pool[2,4] | Relu | Relu | Relu | Conv[3,16,64] |
| 10 | Bn | Bn | Bn | Bn | Relu | Relu | Bn | Bn | Bn | Pool[1,2] |
| 11 | Conv[2,4,64] | Conv[3,16,46] | Conv[1,8,64] | Conv[3,8,64] | Bn | Bn | Conv[2,32,64] | Conv[2,16,64] | Conv[3,4,64] | Relu |
| 12 | Pool[2,4] | Relu | Pool[1,4] | Pool[2,4] | Conv[2,8,64] | Conv[3,4,128] | Pool[1,4] | Relu | Pool[1,4] | Bn |
| 13 | Relu | Bn | Relu | Relu | Relu | Relu | Relu | Bn | Relu | Conv[1,2,128] |
| 14 | Bn | Conv[1,8,64] | Bn | Bn | Bn | Bn | Bn | Conv[1,16,64] | Bn | Pool[1,2] |
| 15 | Conv[3,8,128] | Pool[1,4] | Conv[3,8,64] | Conv[2,2,128] | Conv[1,16,64] | Conv[2,16,128] | Conv[3,2,64] | Pool[1,2] | Conv[3,8,128] | Relu |
| 16 | Relu | Relu | Relu | Pool[1,4] | Pool[1,4] | Pool[1,2] | Relu | Relu | Pool[1,4] | Bn |
| 17 | Bn | Bn | Bn | Relu | Relu | Relu | Bn | Bn | Relu | Fc[512] |
| 18 | Conv[3,32,128] | Conv[3,8,128] | Conv[1,2,64] | Bn | Bn | Bn | Conv[1,2,64] | Conv[2,32,128] | Bn | Relu |
| 19 | Pool[1,4] | Pool[1,4] | Relu | Conv[2,2,128] | Conv[3,4,128] | Conv[1,2,256] | Pool[2,4] | Pool[1,4] | Conv[3,2,256] | Bn |
| 20 | Relu | Relu | Bn | Pool[1,4] | Pool[1,2] | Relu | Relu | Relu | Pool[1,2] | Dropout |
| 21 | Bn | Bn | Conv[2,2,64] | Relu | Relu | Bn | Bn | Bn | Relu | Out |
| 22 | Conv[3,4,256] | Conv[2,2,128] | Pool[2,4] | Bn | Bn | Conv[3,4,256] | Conv[1,8,128] | Conv[2,16,128] | Bn | |
| 23 | Relu | Pool[1,2] | Relu | Conv[1,4,256] | Conv[3,4,256] | Pool[1,2] | Pool[1,1] | Relu | Conv[2,3,512] | |
| 24 | Bn | Relu | Bn | Relu | Relu | Relu | Relu | Bn | Relu | |
| 25 | Conv[3,8,256] | Bn | Conv[2,4,128] | Bn | Bn | Bn | Bn | Conv[1,2,128] | Bn | |
| 26 | Pool[1,2] | Conv[3,2,256] | Relu | Conv[3,2,256] | Conv[3,4,256] | Fc[512] | Fc[512] | Relu | Conv[3,4,512] | |
| 27 | Relu | Pool[1,2] | Bn | Relu | Pool[1,1] | Relu | Relu | Bn | Pool[1,2] | |
| 28 | Bn | Relu | Conv[1,8,128] | Bn | Relu | Bn | Bn | Conv[3,16,128] | Relu | |
| 29 | Fc[512] | Bn | Relu | Conv[2,2,256] | Bn | Dropout | Dropout | Pool[1,4] | Bn | |
| 30 | Relu | Conv[1,8,512] | Bn | Pool[1,2] | Conv[2,4,256] | Out | Out | Relu | Conv[1,3,512] | |
| 31 | Bn | Pool[1,2] | Conv[3,2,128] | Relu | Pool[1,2] | | | Bn | Pool[1,1] | |
| 32 | Dropout | Relu | Pool[1,4] | Bn | Relu | | | Fc[512] | Relu | |
| 33 | Out | Bn | Relu | Fc[512] | Bn | | | Relu | Bn | |
| 34 | | Fc[512] | Bn | Relu | Fc[512] | | | Bn | Fc[512] | |
| 35 | | Relu | Fc[512] | Bn | Relu | | | Dropout | Relu | |
| 36 | | Bn | Relu | Dropout | Bn | | | Out | Bn | |
| 37 | | Dropout | Bn | Out | Dropout | | | | Dropout | |
| 38 | | Out | Dropout | | Out | | | | Out | |
| 39 | | | Out | | | | | | | |

**Architecture Legend**

| Key | Description |
|---|---|
| Conv[X,Y,Z] | Convolutional Layer with Kernel Height X, Kernel Width Y, Z Number of Filters |
| Relu | Rectified Linear Unit Layer |
| Bn | Batch Normalization Layer |
| Pool[X,Y] | Max Pooling Layer with Kernel Height X and Kernel Width Y |
| Fc[X] | Fully Connected Layer with X Number of Units |
| Dropout | Dropout Layer |
| Out | Softmax Classification Layer with 504 Output Units |
